# Interactions across hemispheres in prefrontal cortex reflect global cognitive processing

**DOI:** 10.1101/2025.06.12.659406

**Authors:** Megan E. McDonnell, Akash Umakantha, Ryan C. Williamson, Matthew A. Smith, Byron M. Yu

## Abstract

Brain functions involve processing in local networks as well as modulation from brainwide signals, such as arousal. Dissecting the contributions of populations of neurons to these functions requires knowledge of interactions between brain areas. We investigated these interactions using dual hemisphere recordings of prefrontal cortex in monkeys performing a spatial memory task. To tease apart global processing from local interactions, we applied a novel statistical approach called pCCA-FA (a combination of probabilistic canonical correlation analysis and factor analysis) to analyze trial-to-trial variability in neuronal responses. We found substantial shared variability among neurons within each population, much of which was actually shared across populations and linked to an arousal process. Our work presents a path by which we can leverage multi-area recordings to reveal aspects of brain functions that are hidden in single-area recordings.

## INTRODUCTION

Brain functions such as attention^1,2^, decision making^3^, and working memory^4,5^ rely on coordination among neurons within brain areas as well as across brain areas. Thus, the spiking activity of individual neurons can be influenced by neurons from the same brain area, neurons in other brain areas, global modulatory signals, and more^6^. Teasing apart the impact of each of these influences on neural activity would enable a deeper understanding of how neurons in different brain areas interact to give rise to such brain functions.

To dissect the various influences on neural activity, we can consider how the activity of one neuron covaries with that of other nearby neurons. Many studies have examined how the activity of pairs of neurons in a single brain area covaries from trial-to-trial by computing spike count correlation (*r*_sc_)^7,8^. These pairwise correlations have been widely used as a window to investigate a range of phenomena from cognitive states to circuit architectures^9,10^. To go beyond pairs of neurons, a related body of work has identified the population-wide covariability structure in single brain areas. These population-level approaches have been used to study the trial-to-trial variability underlying sensory encoding^11,12^, attention^13–15^, motor control^16–18^, learning^19–21^, decision-making^22–24^, network structure^25–27^, and more^28^. Taken together, these studies inform our understanding of how neural activity is shared among neurons within a brain area.

Given that most brain functions rely on interactions across brain areas, it is likely that some component of the activity of each brain area is shaped by interactions with other brain areas. Developments in neural recording technology have enabled simultaneous recordings of populations of neurons in multiple brain areas^29–31^, which give rise to the possibility of distinguishing interactions across brain areas from interactions solely within a single brain area^32–35^. Simultaneous recordings from multiple brain areas have been leveraged to study widespread brain functions such as sensory processing^36,37^, motor control^38,39^, learning^40,41^, decision-making^42,43^, arousal^44,45^, and attention^46–49^. However, most statistical methods for analyzing populations of neurons are not designed to distinguish activity shared across areas from activity shared solely among neurons within each area.

Here, we seek to identify components of neural activity that involve interactions across two areas (i.e., an across-area component), as well as interactions solely within each area (i.e., a within-area component). We simultaneously recorded spiking activity from each of the two hemispheres (here referred to as two “areas”) of prefrontal cortex (PFC) in macaque monkeys while they performed a spatial memory task. Utilizing a statistical method we developed called pCCA-FA (a combination of probabilistic canonical correlation analysis and factor analysis), we found substantial shared variance among neurons in PFC, much of which involved neurons across both areas. In fact, the across-area component of PFC activity was larger than the within-area component. We then uncovered a link between the across-area component and a brainwide arousal process. Our work motivates a reinterpretation of shared variance among neurons within a brain area as arising from multiple sources, both within that area as well as via interactions across the brain.

## RESULTS

The coordination among neurons during perception, cognition, and action results in aspects of each neuron’s activity being shared with neurons in the same brain area as well as in other brain areas. Consider the spiking activity of two distinct populations of neurons, recorded simultaneously from two brain areas (area 1 and area 2, Figure 1A). For each neuron, we would like to identify which aspects of its activity are shared with neurons in both areas (pink arrow) and which aspects are shared exclusively with neurons in area 1 (blue arrow). To understand how the activity of area 1, neuron 1 is shared with other neurons (i.e., to quantify the relationships described by the pink and blue arrows in Figure 1A), we would like to partition its trial-to-trial spike count variance into an across-area component (termed “across-area variance”; Figure 1B, pink), a within-area component (termed “within-area variance”; Figure 1B, blue), and a component independent to this neuron (termed “independent variance”; Figure 1B, gray).

**Figure 1.**
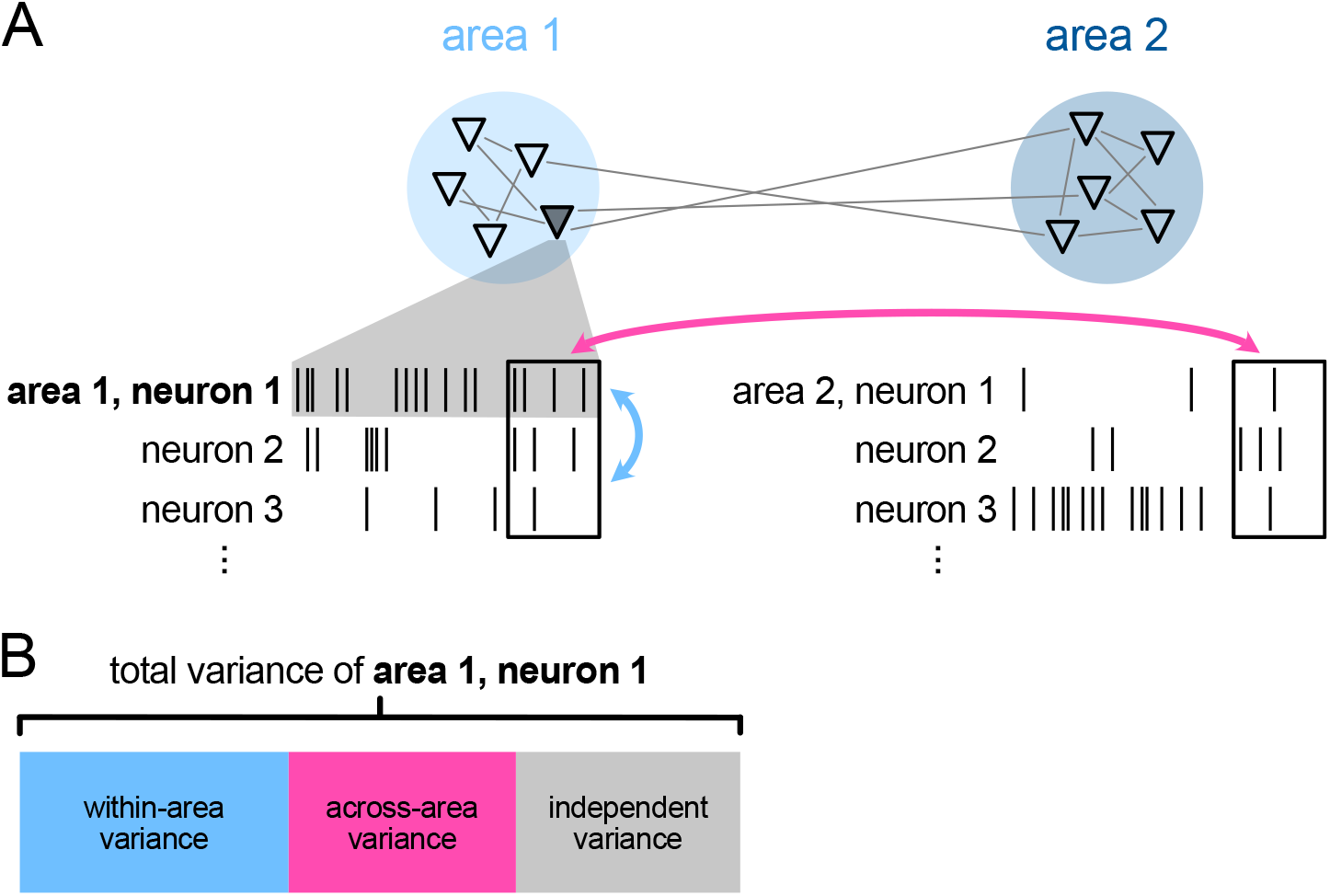
Understanding neuronal variability using spike count variance. (A) Given two populations of neurons, we wish to understand what aspects of area 1, neuron 1’s (shaded triangle) activity (spike counts in boxes) are shared with other neurons across both areas (pink arrow) versus only with other neurons in area 1 (blue arrow). (B) By considering how area 1, neuron 1’s activity covaries with other neurons in the same area or the other area, we can partition its spike count variance into a component that is only shared with neurons in area 1 (“within-area variance”, blue partition), a component that is shared with neurons in both areas (“across-area variance”, pink partition), and a component that is not shared with any other recorded neuron (“independent variance”, gray partition).

### Mean spike count correlation does not adequately partition shared variance

We recorded spiking activity from populations of neurons in each hemisphere of dorsolateral PFC to understand how neural activity was shared either across or within areas (Figure 2A). We began by characterizing the pairwise interactions among neurons within each area using *r*_sc_. The spike counts used to compute *r*_sc_ were taken from a one-second window during the delay period of a memory-guided saccade task (Figure 2B). Throughout this work, we combined trials across conditions after removing condition-specific means, and removed slow-timescale trends to avoid spurious correlations^50^ (see STAR Methods, Figure S1).

**Figure 2.**
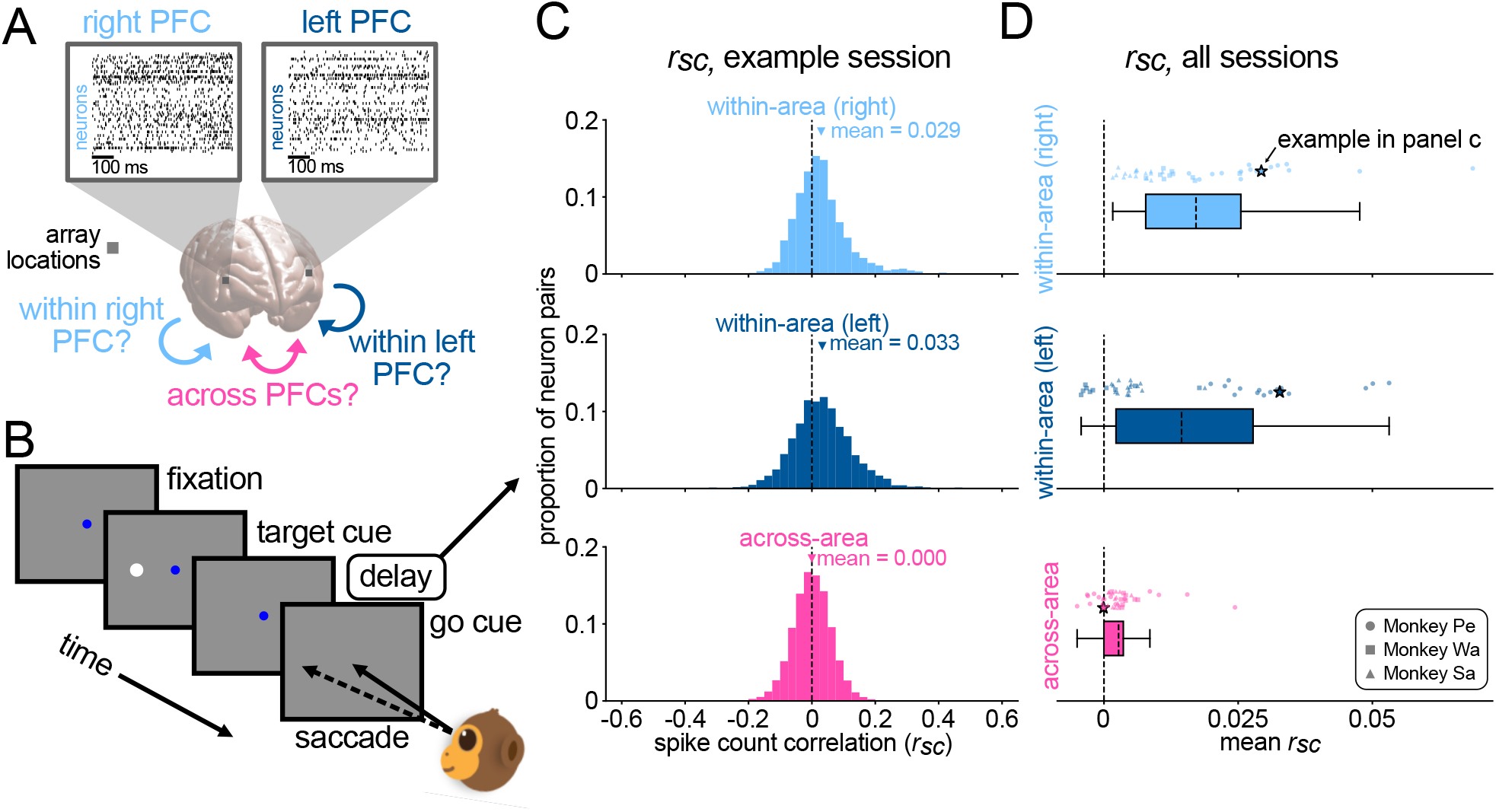
Spike count correlations during a memory-guided saccade task. (A) We recorded from each hemisphere of PFC (referred to here as two brain areas) using multielectrode arrays. The raster plots show spiking activity from each population recorded during an example session. Our goal in this work was to partition neuronal variability in PFC into across- and within-area components. (B) Monkeys performed a memory-guided saccade task in which a brief target cue flash was followed by a variable-length delay period. After the delay period, they were required to report the location of the cue with a saccadic eye movement to the remembered location. The spike counts recorded during the delay period were used for all subsequent analyses. (C) *r*_sc_ distributions for an example session (Monkey Pe). Top, within-area mean *r*_sc_ *>* 0 for pairs of neurons in right PFC (*p <* 0.0001, t-test). Middle, within-area mean *r*_sc_ *>* 0 for pairs of neurons in left PFC (*p <* 0.0001, t-test). Bottom, across-area mean *r*_sc_ ≈ 0 (*p* = 0.55, t-test). (D) Aggregated across sessions, within-area mean *r*_sc_ was greater than across-area mean *r*_sc_ (right PFC: *p <* 0.0001, left PFC: *p <* 0.0001; paired t-test, pooled across animals). Top, within-area mean *r*_sc_ in right PFC. Middle, within-area mean *r*_sc_ in left PFC. Bottom, across-area mean *r*_sc_. Each point is the across- or within-area mean *r*_sc_ from one session. The stars indicate values for the example session in (C). Each box plot covers the first to third quartiles of the mean *r*_sc_ (across- or within-area), with the mean across sessions marked with a line.

The mean *r*_sc_ among pairs of recorded neurons in a brain area is commonly taken to indicate the strength of shared activity among neurons. We computed *r*_sc_ for each pair of neurons within each area (Figure 2C, top and middle, example session), then averaged over the pairs to obtain a within-area mean *r*_sc_ per session (right PFC mean=0.029, left PFC mean=0.033, example session). In the majority of sessions, we found that within-area mean *r*_sc_ values were positive (Figure 2D, top and middle, all sessions).

To understand how activity is shared across areas, one approach is to compute *r*_sc_ between pairs of neurons across areas (i.e., one neuron in each area). Mean *r*_sc_ decreases for pairs of neurons as the physical distance between the neurons increases^51–53^, and has been shown to be smaller for pairs of neurons across versus within brain areas^54^. We observed an across-area mean *r*_sc_ that was close to zero (Figure 2C-D, bottom). This might be taken to suggest that neurons within areas interact more strongly than across areas (Figure 2D, *p <* 0.0001), and that there are little to no interactions across the hemispheres of PFC.

However, there are several limitations to mean *r*_sc_ measurements. First, by averaging over all pairs of neurons (either across or within areas), we ignore the breadth of *r*_sc_ distributions (Figure 2C). For example, if there are pairs of neurons with strong positive correlations and also pairs with strong negative correlations, the mean *r*_sc_ might be close to zero even though many neuron pairs exhibit strong correlations. One way to identify neuron pairs with strong positive and negative correlations is to consider the relative tuning preferences of the neurons^55–58^ (see Figure S2). Second, the within-area mean *r*_sc_ does not reveal whether the correlations shared among neurons in an area are also shared with other areas. Rather, all co-fluctuations among neurons in a brain area are collapsed into the within-area mean *r*_sc_. As such, within-area mean *r*_sc_ could reflect correlations involving interactions with other areas and/or global neuromodulation. Third, mean *r*_sc_ (either across- or within-area) is not designed to identify the many possible population-wide co-fluctuations among neurons. Several distinct changes in population-wide co-fluctuations can result in the same change in mean *r*_sc_ ^59,60^. By collapsing all interactions in one metric, we are unable to separately characterize the many influences on neural activity. Here we desire to leverage the simultaneity of our dual PFC recordings to dissect the multiple population-wide influences on across- and within-area activity (as in Figure 1B).

### pCCA-FA partitions across- and within-area shared variance

To explicitly partition across- and within-area variance, we can leverage statistical methods that take into account all simultaneously-recorded neurons together. In a single brain area, we can identify population-wide co-fluctuation patterns using factor analysis (FA) or related methods^61^. While this provides a more detailed view of population-wide covariability than mean *r*_sc_ ^60^, these methods do not reveal whether activity fluctuations are shared with other brain areas. When recording from two brain areas simultaneously, we can identify variance shared among neurons across areas by employing canonical correlation analysis (CCA) or related methods^32^. However, neither CCA nor its probabilistic variant, pCCA^62^, are designed to also identify variance shared only among neurons within each area (see STAR Methods). Given that none of the aforementioned methods are designed to partition variance into both across-area and within-area components, a new approach is needed.

To address this need, we developed a dimensionality reduction method for two simultaneously recorded brain areas called pCCA-FA (a combination of probabilistic canonical correlation analysis and factor analysis), which partitions the variance of each area’s activity into an across-area component, a within-area component, and a component independent to each neuron. pCCA-FA can be applied to any two populations of simultaneously recorded neurons, and here we focus on the case where each population is a distinct brain area (area 1 and area 2, Figure 3A). We seek to identify latent variables that capture co-fluctuations among neurons across areas (Figure 3A, pink), as well as latent variables specific to each area that capture co-fluctuations among neurons within each area (Figure 3A, light and dark blue for areas 1 and 2, respectively). In total, pCCA-FA identifies three sets of latent variables: one across-area set, and a separate within-area set for each area. Because the within-area latent variables are specific to each area, the number of within-area latent variables for each area need not be the same.

**Figure 3.**
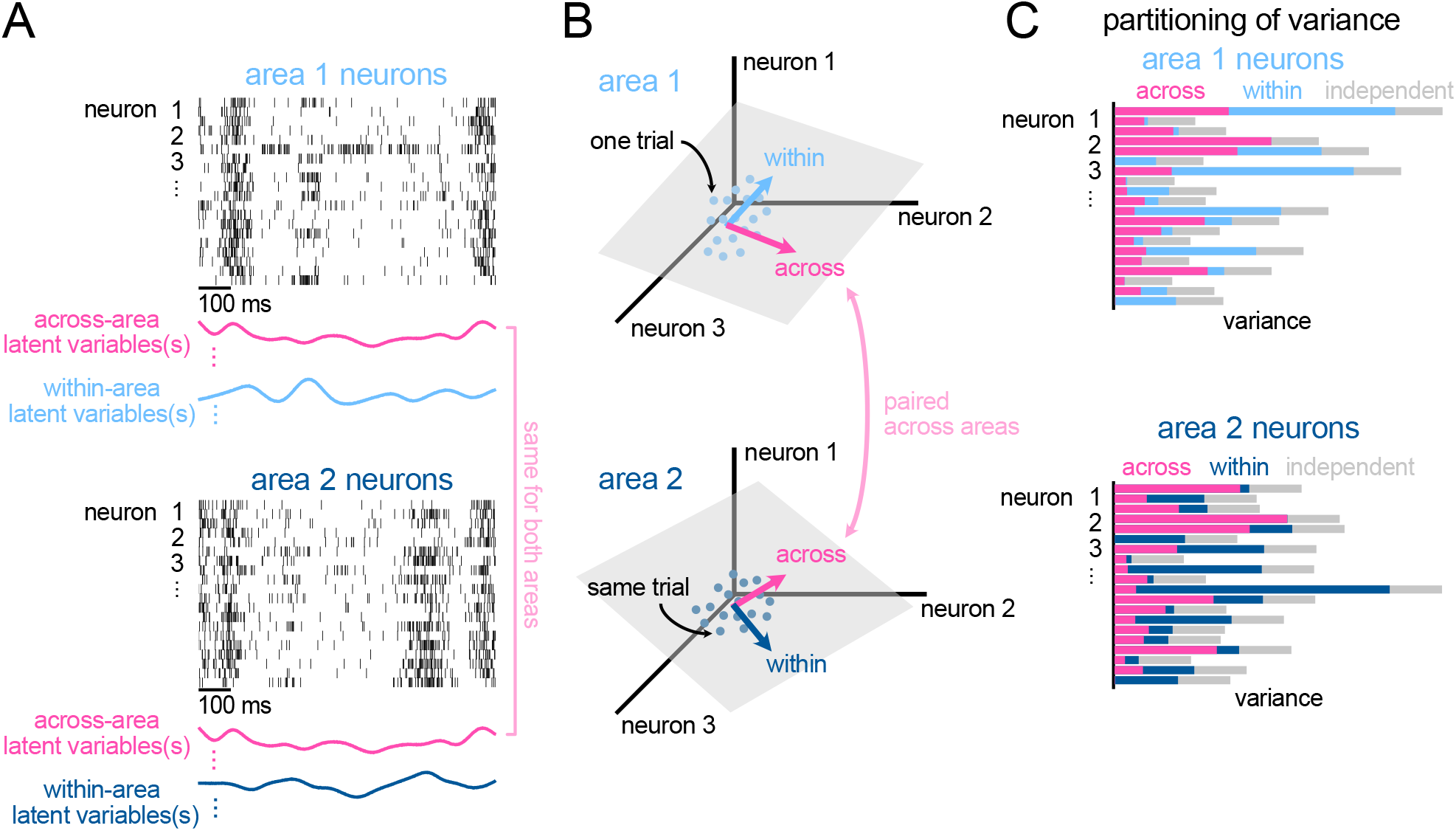
pCCA-FA partitions across- and within-area shared variance. (A) Simulated neural population activity from two brain areas (illustrated here using 20 neurons in each brain area) share an across-area latent variable (pink trace, same for both areas). Each area has its own within-area latent variable (light and dark blue trace for area 1 and area 2, respectively). The activity of each neuron can be constructed as a weighted linear combination of these latent variables. (B) We define a population activity space for each area, where each coordinate axis represents the activity of one neuron. Each point represents the spike counts on a particular trial. For each point in area 1’s population activity space, there is a corresponding point in area 2’s population activity space for the same trial. Each latent variable can be represented by a co-fluctuation pattern or vector in the corresponding population activity space, which maps the latent variable to the neurons’ activity. Across-area co-fluctuation patterns map the across-area latent variables to neurons in both areas (pink arrows). As there is a single set of across-area latent variables that is common to both areas, the across-area co-fluctuation patterns are paired across areas. Within-area co-fluctuation patterns map the within-area latent variables to the neurons in the corresponding area (light and dark blue arrows for area 1 and 2, respectively). There is a separate set of within-area latent variables for each area, so the within-area co-fluctuation patterns are not paired. (C) pCCA-FA partitions each neuron’s variance into three components: across-area variance (pink), within-area variance (light and dark blue for area 1 and area 2 respectively), and independent variance (gray). Each row of the rasters in (A) and each coordinate axis in (B) is represented as a row in (C). The relative size of each partition indicates the proportion of each neuron’s variance that is attributed to the across(pink) or within-area (blue) component.

Each latent variable represents a characteristic co-fluctuation pattern among the neurons. A co-fluctuation pattern can be represented as a vector in the corresponding population activity space, where each coordinate axis represents the activity of one neuron in that brain area (Figure 3B). Each across-area latent variable identified by pCCA-FA is represented by one co-fluctuation pattern in each area (Figure 3B, pink). In contrast, each within-area latent variable is represented by one co-fluctuation pattern in either area 1 or 2 (Figure 3B, light and dark blue, respectively). For each neuron, we can quantify the proportion of its spike count variance explained by each set of latent variables. pCCA-FA partitions each neuron’s variance into a component that is described by the across-area latent variables (termed “across-area variance”, Figure 3C, pink), and a component that is described by that area’s within-area latent variables (termed “within-area variance”, Figure 3C, light and dark blue for area 1 and 2, respectively). The residual variance not accounted for by across- or within-area latent variables is independent to each neuron (termed “independent variance”, Figure 3C, gray). Thus, pCCA-FA explicitly partitions the spike count variance of each neuron into across-area, within-area, and independent components (as in Figure 1B).

To summarize the outputs of pCCA-FA, we define the following two metrics. First, to understand the extent to which a particular set of latent variables explains a neuron’s spike count variance, we define the across- and within-area percentage of variance shared among neurons (%sv). The values of this metric reflect the relative size of the bars in Figure 3C. Across-area %sv is defined as the proportion of spike count variance explained by the across-area latent variables, averaged across neurons in each brain area (see Equation 8). Within-area %sv is defined analogously using the within-area latent variables (see Equation 9). Thus, each brain area has both an across-area %sv and a within-area %sv. Second, we quantify the complexity of the interactions described by each set of latent variables by computing its shared dimensionality (*d*_*shared*_)^26,63^. Across- and within-area *d*_*shared*_ are defined as the number of latent variables required to explain the across- or within-area variance, respectively (see STAR Methods).

Before applying pCCA-FA to neural recordings, we performed extensive model validation. We verified that pCCA-FA accurately identified the ground truth %sv and *d*_*shared*_ using simulated data (Figure S3A-B and Figure S4), and found that it required fewer trials to do so than pCCA (Figure S3C-D). Further, pCCA-FA outperformed other dimensionality reduction techniques (pCCA and FA) on neural recordings (Figure S5). Thus, the pCCA-FA framework successfully captures how neurons in two brain areas covary by partitioning variance into across- and within-area components.

### Across-area interactions were substantial and often greater than within-area interactions

To identify across- and within-area variance in our PFC recordings, we fit pCCA-FA to the same neural activity (with the same preprocessing) as in the *r*_sc_ analyses (Figure 2). For each session, pCCA-FA estimated a single set of across-area latent variables common to both areas and a separate set of within-area latent variables for each area. From the mean *r*_sc_ values in Figure 2, one might expect to uncover a large within-area component and small across-area component using pCCA-FA.

To determine if this was the case, we computed across- and within-area %sv and *d*_*shared*_. For the same example session as in Figure 2C, we found that a substantial portion of variance was shared across areas (14.97% and 17.47%, for right and left PFC, respectively), compared to the variance shared exclusively within each area (5.65% and 7.47%, for right and left PFC, respectively) (Figure 4A). This trend was evident across sessions, with the across-area %sv being larger than the within-area %sv (Figure 4B, *p <* 0.0001, pooled across animals). Thus, substantial across-area interactions were present in the activity and identified by pCCA-FA, contrary to the intuition from measuring pairwise across-area *r*_sc_. This discrepancy can be understood by considering the sign of pairwise correlations among neurons (Figure S2). Mean *r*_sc_ averages over positive and negative correlations found in individual pairs of neurons, whereas %sv accounts for both positive and negative correlations among neurons.

**Figure 4.**
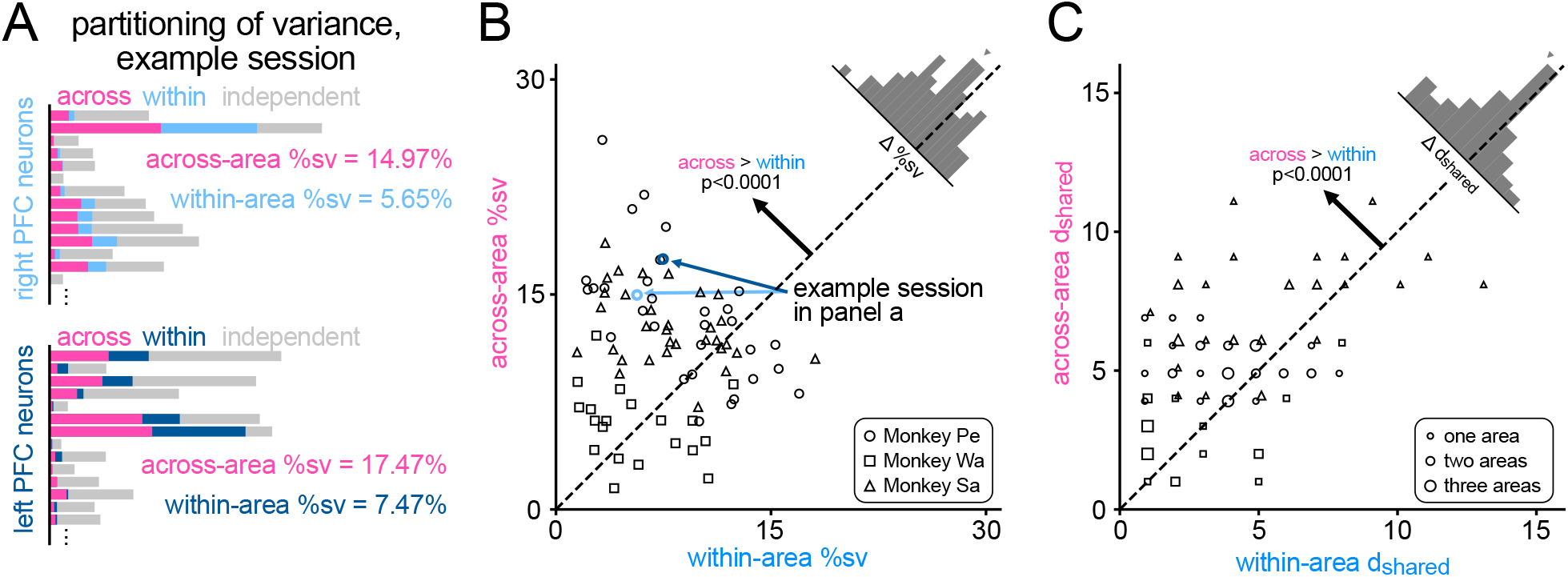
Across-area interactions were substantial and often greater than within-area interactions. (A) pCCA-FA partitions the variance of dual-hemisphere PFC recordings (same example session as Figure 2B, Monkey Pe). Same conventions as in Figure 3C. Across-area %sv was 14.97% and 17.47%, and within-area %sv was 5.65% and 7.47%, for right and left PFC, respectively. For visual clarity, only 14*/*79 neurons in right PFC and 14*/*83 neurons in left PFC are shown. Note, though the across-area latent variables are common to both areas, the across-area %sv can have a unique value, depending on the individual variances of each neuron. (B) Aggregated across sessions, across-area %sv (mean 11.50%) was larger than within-area %sv (mean 7.66%; *p <* 0.0001 pooled across animals; Monkey Pe: *p* = 0.00048; Monkey Wa: *p* = 0.40, Monkey Sa: *p <* 0.0001; paired t-test). Each session contributes two points, one each for right and left PFC. The gray histogram shows the difference between across- and within-area %sv for each session (Δ = within-area %sv - across-area %sv), with the mean difference indicated by a triangle. The example session in (A) is marked in light and dark blue for right and left PFC, respectively. (C) Aggregated across sessions, across-area *d*_*shared*_ (mean 5.41) was larger than within-area *d*_*shared*_ (mean 4.06; *p <* 0.0001 pooled across animals; Monkey Pe: *p* = 0.0013, Monkey Wa: *p* = 0.32, Monkey Sa: *p <* 0.0001; paired t-test). Same conventions as in (B). Each session contributes two values, one for each area. Because *d*_*shared*_ is integer-valued, the values from multiple sessions overlap. The size of each symbol indicates the number of areas across sessions with the particular values of *d*_*shared*_ (larger symbol indicates more sessions). Symbols for each monkey are jittered slightly for visual clarity.

Having found that neurons tended to more strongly covary across areas than within each area, we next considered the complexity of these interactions. We found that the across-area *d*_*shared*_ was also larger than the within-area *d*_*shared*_ (Figure 4C, *p <* 0.0001, pooled across animals). This indicated that, in addition to being stronger, the interactions we identified across areas were more complex than the interactions within each area.

### Across- and within-area components involved different co-fluctuation patterns

We next sought to understand the population-wide co-fluctuation patterns represented by each set of latent variables, as they provide insight into the nature of the co-fluctuations among neurons (Figure 5A). We first asked whether the across- or within-area co-fluctuation patterns involved the entire populations increasing or decreasing their activity together, which might be the case for a global change in responsivity. For this analysis, we focused on the first co-fluctuation pattern of each ordered set, as it reflected a large proportion of the across- or within-area variance (Figure S6A-B). We found that global changes in responsivity, and thus population-wide increases and decreases in activity, were not consistent with the across- or within-area cofluctuation patterns we identified (Figure 5B, Figure S6C).

**Figure 5.**
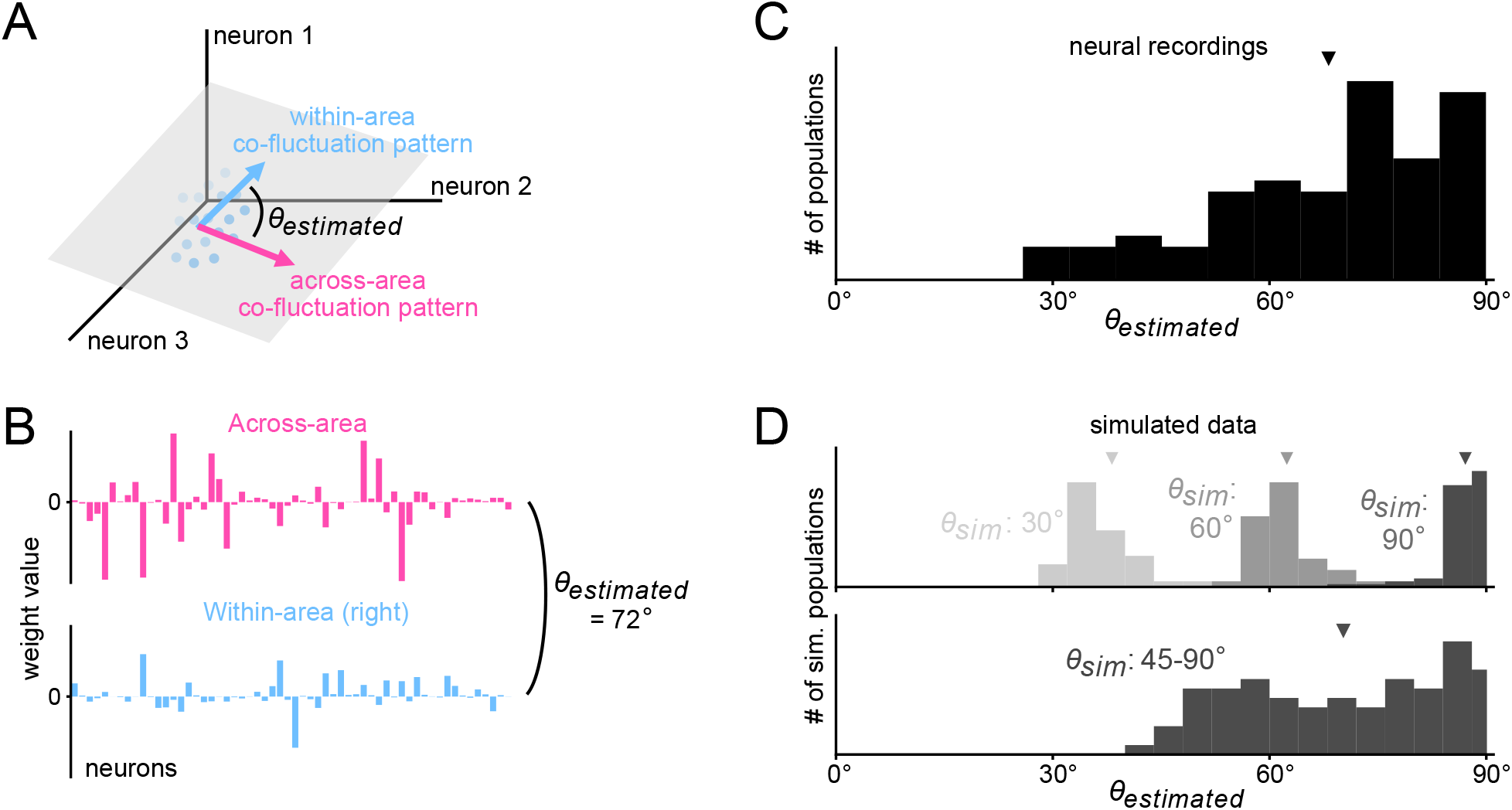
Across- and within-area components involved different co-fluctuat patterns. (A) The across- (pink) and within-area (blue) co-fluctuation patterns identified by pCCA-FA can be represented as vectors in each population activity space. To determine how similar the top across- and within-area co-fluctuation patterns were, we computed the angle between them (*θ*_*estimated*_). (B) Weights of the top across- (pink) and within-area (blue) co-fluctuation patterns from an example session (Monkey Sa, right PFC). The height of each bar indicates the weight of one neuron for that co-fluctuation pattern. For this example session, *θ*_*estimated*_ = 72^°^. (C) Angles between the top across- and within-area co-fluctuation patterns (*θ*_*estimated*_) in neural recordings for all sessions (mean *θ*_*estimated*_ = 68.2^°^ marked with a triangle). Each session contributed one angle per brain area (i.e., two angles per session). (D) Simulations in which the ground truth angle between the top across- and within-area cofluctuation patterns was specified (*θ*_*sim*_, see STAR Methods) were performed to generate chance distributions. Same conventions as in (C). Top, three simulations were performed in which *θ*_*sim*_ was specified as a constant value across all simulated datasets (*θ*_*sim*_ ∈ 30, 60, 90^°^). For each value of *θ*_*sim*_, one simulated dataset was generated for each session in (C). The number of neurons, co-fluctuation patterns, and trials were matched to the neural recordings, as well as the amount of shared variance (i.e., the scaling of each co-fluctuation pattern). The *θ*_*estimated*_ from neural recordings (C) was greater than the *θ*_*estimated*_ from simulated data for *θ*_*sim*_ = 30^°^ (*p <* 0.0001, paired t-test) and for *θ*_*sim*_ = 60^°^ (*p* = 0.00092, paired t-test), but not for *θ*_*sim*_ = 90^°^ (*p* = 1.0, paired t-test). Bottom, an additional simulation was performed in which the value of *θ*_*sim*_ was drawn uniformly and independently for each dataset between 45 and 90°. The *θ*_*estimated*_ from neural recordings (C) was not statistically different from the *θ*_*estimated*_ from simulated data (*p* = 0.40, paired t-test).

We next asked whether across- and within-area co-fluctuation patterns were similar to each other by computing the angle between them in population activity space (*θ*_*estimated*_; Figure 5A). They might be similar if two different processes (one global and one local) had a similar effect among the neurons. Alternatively, each process might engage distinct circuit mechanisms, yielding different co-fluctuation patterns. In an example session, we found that the across- and within-area co-fluctuation patterns appeared visually distinct (Figure 5B), and produced a *θ*_*estimated*_ of 72^°^. In all sessions, we found that the across- and within-area co-fluctuation patterns had large *θ*_*estimated*_ (Figure 5C), often close to 90^°^, which would indicate distinct co-fluctuation patterns.

To assess the extent to which the measured *θ*_*estimated*_ reflected true differences in the across- and within-area co-fluctuation patterns, we created a simulation in which we defined the ground truth angle between across- and within-area co-fluctuation patterns (*θ*_*sim*_, see STAR Methods, Figure S4). We simulated activity in each population with a specified value of *θ*_*sim*_ that was otherwise matched to the neural recordings (see STAR Methods). We found that, for any given *θ*_*sim*_, the resulting distributions of *θ*_*estimated*_ (Figure 5D, top) were narrower than what we observed in the neural recordings (Figure 5C). Instead, if we sampled a different value of *θ*_*sim*_ for each simulated dataset uniformly from 45 − 90^°^, we recovered a distribution of *θ*_*estimated*_ (Figure 5D, bottom) that was similar to that recovered from the neural recordings (*p* = 0.40). Together, these results indicated that across- and within-area components involved different co-fluctuation patterns that varied in their similarity from session to session.

### Across-area latent variables were related to pupil diameter

Given that we found substantial across-area variance (Figure 4) involving different co-fluctuation patterns than the within-area component (Figure 5), we considered whether it might be related to a brainwide process. Arousal is known to modulate activity of neurons throughout the brain^44,45,64–66^. In addition, arousal state has often been linked to changes in the size of the pupil^67,68^. Trial-by-trial and faster-timescale evoked changes in pupil diameter have been linked to arousal-related changes in neural activity in both cortical^69–72^ and subcortical^67,73–75^ regions. Given that arousal has been found to be a brainwide phenomenon, we wondered if it was related to the across-area variance we observed. We therefore asked how well the latent variables identified by pCCA-FA explained changes in pupil diameter (Figure 6A).

**Figure 6.**
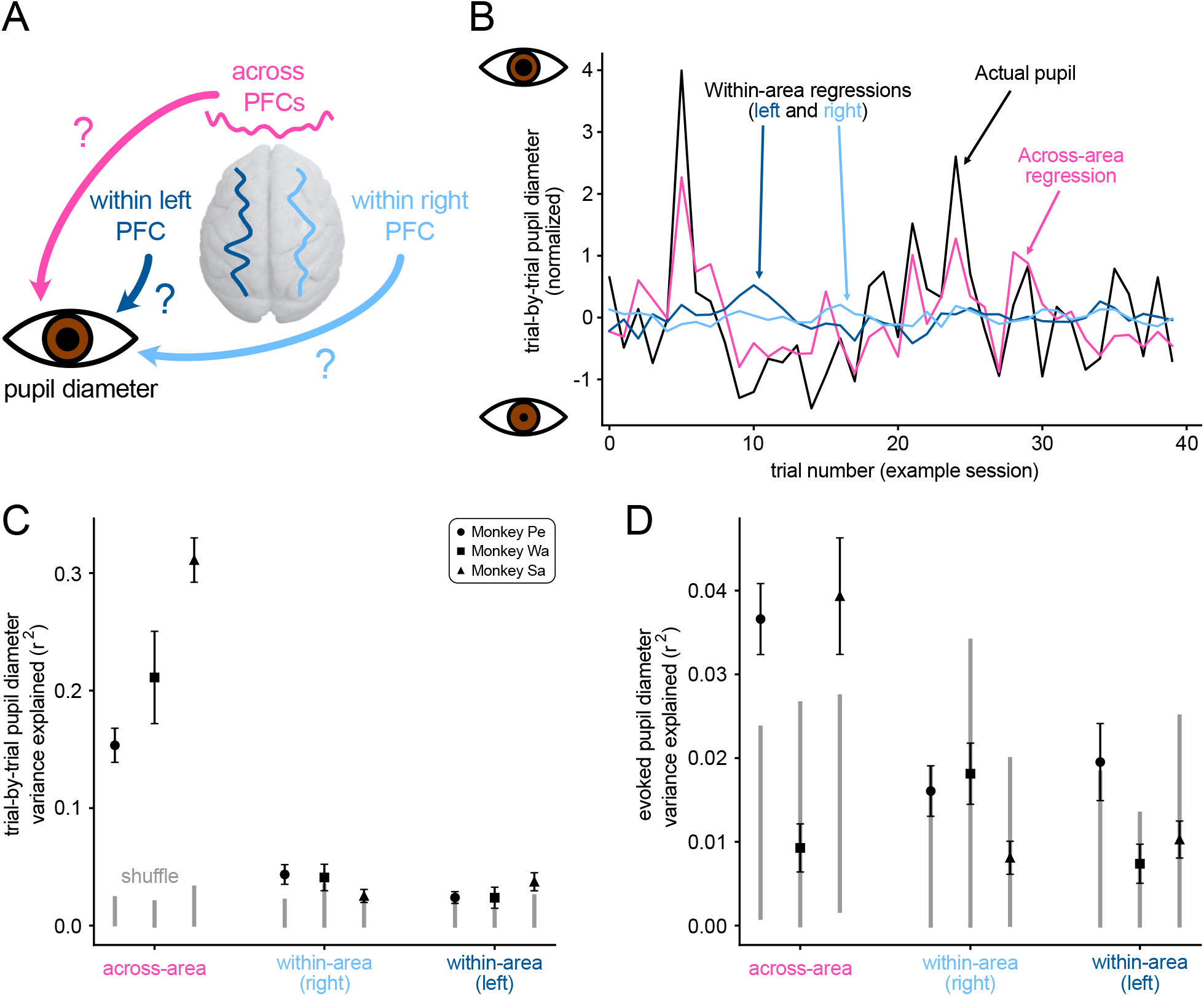
Across-area latent variables were related to pupil diameter. (A) We examined whether across- (pink) or within-area (light and dark blue for right and left PFC, respectively) latent variables explained variance in the pupil diameter. (B) Fluctuations in pupil diameter (black, one normalized value per trial, see STAR Methods) for an example session (Monkey Sa). For visual clarity, only 40 trials are shown. Regression onto the pupil diameter using across-area latent variables (pink) and within-area latent variables identified by pCCA-FA (light and dark blue for right and left PFC, respectively) are overlaid. For this session, across-area latent variables explained more trial-to-trial variance in the pupil diameter (*r*^2^ = 0.35) than within-area latent variables (right PFC: *r*^2^ = 0.013, left PFC: *r*^2^ = 0.052). (C) Aggregated across sessions, *r*^2^ values for trial-by-trial pupil diameter were greater for across-area latent variables (mean *r*^2^ = 0.23) than for within-area latent variables (right PFC, mean *r*^2^ = 0.036; left PFC, mean *r*^2^ = 0.029) (across *>* within right PFC, *p <* 0.0001; across *>* within left PFC, *p <* 0.0001; pooled across animals; Monkey Pe, across *>* within right PFC, *p <* 0.0001; Monkey Pe, across *>* within left PFC, *p <* 0.0001; Monkey Wa, across *>* within right PFC, *p* = 0.0023; Monkey Wa, across *>* within left PFC, *p* = 0.00075; Monkey Sa, across *>* within right PFC, *p <* 0.0001; Monkey Sa, across *>* within left PFC, *p <* 0.0001; paired t-test). Gray bars indicate 95% confidence intervals of the shuffle distribution, obtained by regressing onto the trial-by-trial pupil diameter on session *j* using the latent variables on session *i*, where *i* ≠ *j*. We repeated the same analysis but used only the top co-fluctuation pattern of each type for the regression model to control for differences in across- and within-area dimensionality, and found similar results (right PFC: *p <* 0.0001, left PFC: *p <* 0.0001, pooled across animals; paired t-test). (D) Aggregated across sessions, *r*^2^ values for explaining evoked pupil diameter aggregated across sessions were greater for across-area latent variables (mean *r*^2^ = 0.031) than for within-area latent variables (right PFC, mean *r*^2^ = 0.014; left PFC, mean *r*^2^ = 0.013) for two of three animals (across *>* within right PFC, *p* = 0.00010; across *>* within left PFC, *p <* 0.0001; pooled across animals; Monkey Pe, across *>* within right PFC, *p* = 0.00019; Monkey Pe, across *>* within left PFC, *p* = 0.011; Monkey Wa, across *>* within right PFC, *p* = 0.94; Monkey Wa, across *>* within left PFC, *p* = 0.33; Monkey Sa, across *>* within right PFC, *p* = 0.00059; Monkey Sa, across *>* within left PFC, *p* = 0.00076; paired t-test). When controlling for the number of latent variables, we found similar results. Same conventions as in (C).

To answer this question, we measured the mean pupil diameter during the delay period and adjusted the pupil diameter values values to remove slow fluctuations in an analogous manner to the neural activity (see STAR Methods). We used the across- or within-area latent variables identified by pCCA-FA (i.e., either *d* across-area latent variables or *d*_*m*_ within-area latent variables for area *m*; see STAR Methods) to compute a trial-by-trial regression of the pupil diameter for each set of latent variables (Figure 6B). Across sessions, we found that the across-area latent variables explained substantially more variance in the pupil diameter than the within-area latent variables (right PFC, *p <* 0.0001; left PFC, *p <* 0.0001; Figure 6C).

Next, we asked whether the across- and within-area latent variables could explain variance in the pupillary evoked response magnitude, defined as the fast-timescale change in pupil diameter just after visual stimulus presentation (see STAR Methods). We found that the across-area latent variables also explained more variance in the pupillary evoked response than the within-area latent variables (right PFC, *p* = 0.0001; left PFC, *p <* 0.0001; Figure 6D). Thus, the across-area latent variables were able to explain multiple aspects of pupil-linked arousal. Taken together, these results are consistent with a link between the coordinated activity across brain areas and global cognitive processes such as arousal.

## DISCUSSION

In this work, we asked to what extent neural activity is shared across versus within hemispheres of PFC by considering how the spike count variance of the two brain areas covaried from trialto-trial. Applying a novel statistical approach called pCCA-FA, we found that of the substantial shared variance among PFC neurons, a large proportion involved neurons in both areas. The component of neural activity shared among neurons in both areas (termed “across-area variance”) was larger in magnitude (%sv) and complexity (*d*_*shared*_) than the component shared only among neurons in one area (termed “within-area variance”), in contrast to what was suggested by pairwise *r*_sc_ analyses. Additionally, we found that the across-area component was related to pupil diameter. Our approach thus leveraged multi-area recordings in PFC to uncover signatures of global cognitive processing that were otherwise hidden in single-area analyses.

Spike count variance shared across hemispheres of PFC may arise from a variety of sources. First, it may come from brainwide circuits involved in neuromodulation. One such pathway involves norepinephrine signaling from the locus coeruleus^76,77^, which has both projections to PFC^78,79^ and influence on pupil diameter^80^. Correlations have been observed between pupil diameter and neural activity across the brain^36,39,45^, supporting the notion that this may indeed arise from a global source. Second, across-area variance may arise from direct communication between the hemispheres of PFC, where previous work has demonstrated anatomical^81,82^ as well as functional^83^ connections. Third, across-area variance may result from shared signals about the external world, arising from sensory inputs and common motor plans^36,39^.

Although we emphasized the variance shared across areas, we also identified a substantial amount of variance that was shared only among the neurons within each area. We labeled this variance “within-area”, but we note that it could be shared with other non-recorded brain areas. The origin of what we termed within-area variance may lie in a number of possible sources. First, representations of visual fields in PFC are topographically arranged with a contralateral bias, resulting in each hemisphere of PFC encoding different, although partially overlapping, information^84–86^. Second, internal states like spatial attention tie neurons together in one part of the visual field. For example, previous studies of visual area V4 during spatial attention tasks have identified attention-related components of neural activity that are hemisphere-specific^14,46^. This type of selective modulation may account for some of the within-area variance we report here. Third, within-area variance may result from constraints on patterns of neural activity imposed by the cortical circuitry (e.g., horizontal connections) in each hemisphere^87,88^. In network models, spatial clustering architecture that may be specific to local circuits within a hemisphere can have a profound influence on trial-to-trial variability^25–27,89,90^.

With the identification of within-area variance, pCCA-FA allowed us to isolate components of neural activity that were not shared between the two hemispheres of PFC. Previous work has described a gating of signals across areas, resulting in a communication subspace^91^. pCCA-FA identified what would be termed a communication subspace between the two hemispheres of PFC: we found distinct across- and within-area co-fluctuation patterns, indicating that not all signals were shared between areas. Put another way, the existence of the within-area latent variables in our PFC recordings indicated that a communication subspace exists between the hemispheres.

pCCA-FA improves upon CCA and its probabilistic variant (pCCA) in identifying across-area interactions. First, CCA and pCCA are designed only to identify across-area variance and do not partition the remaining variance into a component that is shared among neurons within each area from variance that is independent to each neuron. Thus, pCCA-FA provides a more thorough dissection of the population activity in two brain areas. Second, even in cases where only across-area variance is of interest, pCCA-FA produced a more accurate estimate of across-area variance, especially in the limited data regime (i.e., low trial counts), than pCCA (Figure S3C- D). This was due to the pCCA-FA model having a smaller number of parameters than pCCA (see Equations 2 and 4). The tradeoff is the increased computation time of pCCA-FA compared with pCCA for jointly optimizing over the hyperparameters of across- and within-area dimensionality. Part of this cost can be mitigated by employing efficient hyperparameter optimization techniques^92,93^. For these reasons, we advocate for the use of pCCA-FA in place of CCA or pCCA for studying interactions between brain areas.

pCCA-FA fills an unmet need in the set of exploratory tools tailored for studying the trial-to-trial variability of neural activity. When studying neurons from a single brain area, FA is among the simplest approaches for separating variance that is shared among neurons from independent variance. For two brain areas, pCCA-FA is among the simplest methods for separating variance that is shared across areas from variance that is only shared among neurons within each area. A related body of work has incorporated time series for studying moment-to-moment fluctuations of neural activity within each trial, such as Gaussian Process Factor Analysis^94^ and related methods^95^ for single brain areas or Delayed Latents Across Groups^93^ and related methods^34,96–100^ for multiple brain areas. As multi-area population recordings continue to increase in prevalence^29–31^, pCCA-FA and related methods will become increasingly essential for dissecting the rich structure of across- and within-area interactions.

## RESOURCE AVAILABILITY

### Lead contact

Requests for further information and resources should be directed to and will be fulfilled by the lead contact, Byron M. Yu.

### Materials availability

This study did not generate new materials.

## Data and code availability

1. Code for fitting and using the pCCA-FA model will be available as of the date of publication.
2. Data and analysis code used in this paper will be available as of the date of publication.

## ACKNOWLEDGMENTS

The authors would like to thank Samantha Schmitt for help with animal training and data collection, our animal care staff, and Evren Gokcen, Emilio Salazar-Gatzimas, and Sam Snyder for helpful discussions. M.E.M. was supported by NIH T32 EB029365. B.M.Y. and M.A.S. were supported by NIH R01 MH118929, NSF NCS BCS 1734916/1954107, NSF NCS DRL 2124066/2123911, and NIH R01 EB026953. B.M.Y was supported by NIH RF1 NS127107, NIH R01 EY035896, and Simons Foundation 543065 and NC-GB-CULM-00003241-05. M.A.S. was supported by NIH R01 EY029250 and NIH R01 MH128393.

## AUTHOR CONTRIBUTIONS

Conceptualization, M.E.M., A.U., R.C.W., M.A.S., B.M.Y.; Data curation: M.E.M., A.U., R.C.W.; Formal analysis, M.E.M., A.U.; Funding acquisition, M.A.S., B.M.Y.; Investigation, M.E.M., A.U., R.C.W.; Methodology, M.E.M., A.U., R.C.W., M.A.S., B.M.Y.; Software, M.E.M., A.U.; Resources, M.A.S., B.M.Y.; Supervision, M.A.S., B.M.Y.; Validation, M.E.M., A.U.; Visualization, M.E.M., A.U., R.C.W.; Writing – original draft, M.E.M., A.U., R.C.W.; Writing – review & editing, M.E.M., A.U., R.C.W., M.A.S., B.M.Y.

## SUPPLEMENTAL INFORMATION INDEX

Figures S1-S6 and their legends in a PDF

## STAR METHODS

### Key resources table

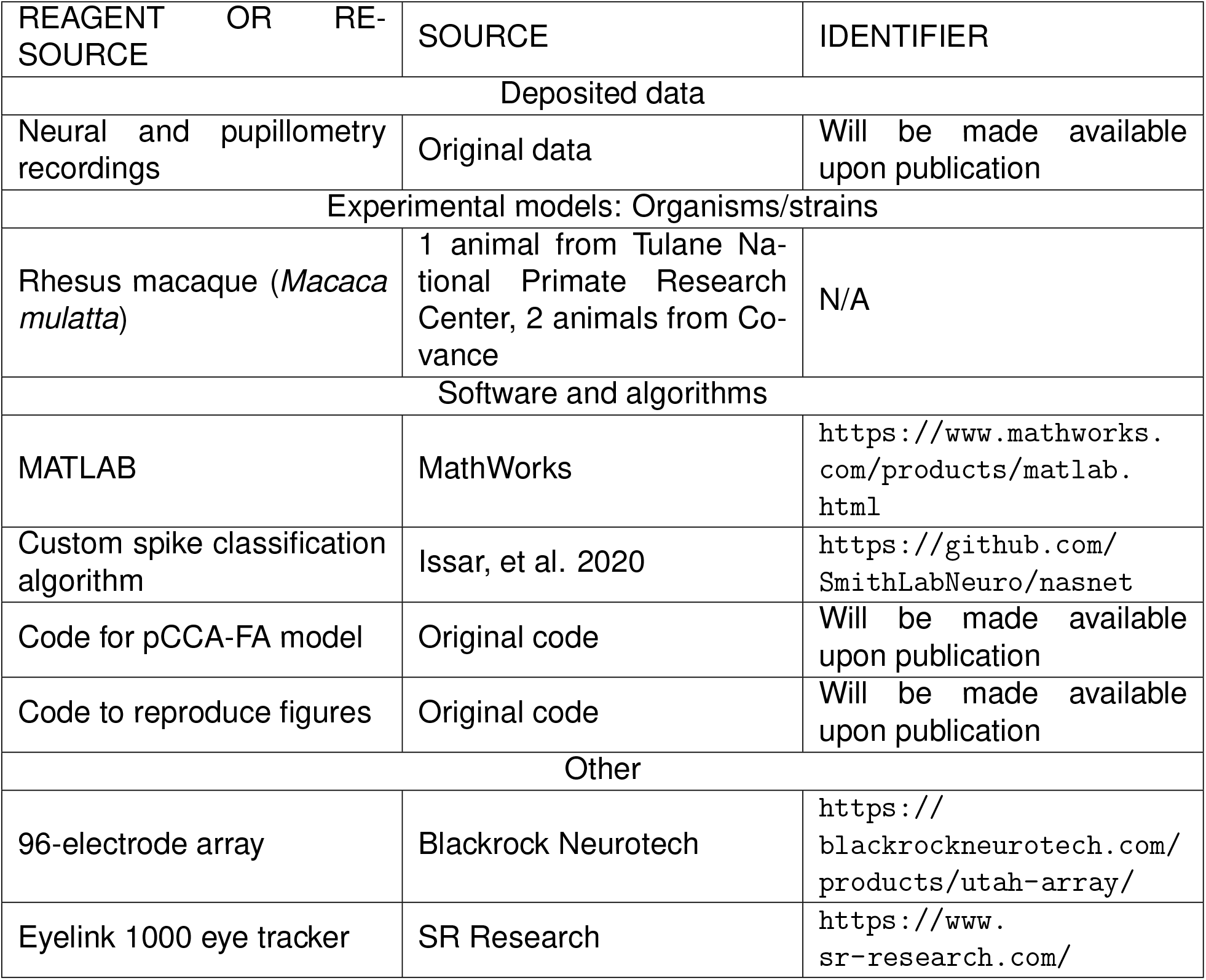

### Experimental model and study participant details

#### Experimental model

Three male rhesus macaque monkeys (*Macaca mulatta*) were used in this study. They were 10 (Monkey Pe), 7 (Monkey Wa), and 7 (Monkey Sa) years old at the time of data collection. The animals were housed singly in a room operating on a standard 12 hour light/dark cycle and were provided with an enhanced enrichment program. All experimental procedures were approved by the Institutional Animal Care and Use Committees of Carnegie Mellon University and the University of Pittsburgh, and were in accordance with the United States National Research Council’s Guide for the Care and Use of Laboratory Animals.

### Method details

#### Surgical preparation

Each animal was surgically implanted with a titanium headpost, which was fixed onto the skull using titanium screws. This was necessary to limit head movement such that the position of the eyes could be tracked and neural activity recorded. Animals were then each implanted with two 96-electrode “Utah” arrays (Blackrock Neurotech, Salt Lake City, UT). Electrode arrays were placed on the prearcuate gyrus just anterior to the arcuate sulcus, one in each hemisphere. Surgeries were performed in sterile conditions under general anesthesia using isoflurane.

#### Electrophysiological methods

Animals were positioned 36 cm from a 21” cathode ray tube monitor with 1024×768-pixel resolution and a refresh rate of 100 Hz. Task stimuli were generated using custom-designed software in Matlab (Mathworks, Natick, MA) that utilized the Psychophysics toolbox extensions^101–103^. Signals from the implanted electrodes were band-pass filtered (0.3 - 7500 Hz) and then digitized at 30,000 Hz. For each electrode, spiking waveforms were defined as a 52-sample (1.73 ms) window of the filtered voltage signal triggered by the signal crossing a predefined threshold, and stored for offline analysis. The threshold was defined as a multiple of the root-mean-square voltage of a brief epoch of the raw signal on each electrode collected at the beginning of each session. All behavioral data and neural activity were recorded using a Grapevine recording system (Ripple, Salt Lake City, UT) for further offline processing and analysis.

#### Behavioral Task

Animals were trained to perform a standard memory-guided saccade task^104^. At the beginning of each trial, a 0.5 degree diameter blue fixation circle appeared at the center of a gray screen. The animal initiated fixation within an invisible window (Monkey Pe: 2.4, Monkey Wa: 1.8, Monkey Sa: 2.9 degrees radius) centered on the fixation spot and then 200 ms later a white target flashed in the animal’s periphery (Monkey Pe: 11.8 or 17.5 degrees from fixation depending on the session, Monkey Wa: 11.6 degrees from fixation, Monkey Sa: a single value in the range of 8.0 - 15.6 degrees from fixation depending on the session, with one session containing multiple randomly interleaved distances in this range) at one of 4, 8, or 16 locations depending on the session. The white target remained on the screen for either 100, 200, or 400 ms (depending on the session) after which it was removed from the screen. The animal maintained fixation for a delay period drawn randomly from a uniform distribution between 1.5 and 3 seconds. After the delay, the fixation spot disappeared, which served as the go cue for the animal to initiate a saccade toward the remembered location where the white target flash had appeared. The animal had 500 ms (Monkey Sa: 400 ms) to initiate a saccade, and an additional 200 ms to reach the target location. Successful completion of a saccade was defined by the animal maintaining fixation within an invisible window (Monkey Pe: 4.1, Monkey Wa: 5.9 or 7.0, Monkey Sa: 4.1 degrees radius) centered at the target location for at least 100 ms. A liquid reward was delivered for a saccade to the correct location. For a subset of sessions, a dim white circle was flashed at the target location after saccade initiation to assist the animal in target acquisition. Trials were pseudo-randomized in blocks during which the animal was required to correctly complete all target directions before beginning a new block. In total, we collected 42 sessions of data (16 in Monkey Pe, 10 in Monkey Wa, and 16 in Monkey Sa). The number of trials per session for Monkey Pe was 729.8 ± 173.4 (mean ± standard deviation), for Monkey Wa was 463.5 ± 170.1, and for Monkey Sa was 850.1 ± 284.2.

#### Preprocessing of neural activity

We used a neural network described previously^105^ to classify waveforms as spikes or noise. Briefly, the network was trained on human-sorted waveforms to distinguish between waveforms putatively of neural origin and waveforms not of neural origin. Using this network, we removed threshold crossings that were unlikely to be of neuronal origin, and called the remaining waveforms “spikes”. We further removed channels that had very low firing rates or very high trialto-trial variability. To do this, we first binned neural activity by counting spikes during the delay period between target offset and saccade initiation cue. We then removed channels with mean spike count lower than 2 spikes/second or Fano factor greater than 10.

We also removed channels affected by artifactual cross-talk due to electrical coupling. For each pair of channels, we flagged spikes as coincident if they occurred within 100 *µ*s of each other. If either channel in the pair had 20% or more of its spikes flagged as coincident, we flagged that pair as having artifactual crosstalk. This procedure provided an index for each channel of how many other channels it was coincident with. We then iteratively removed the channel coincident with the highest number of other channels until no crosstalk remained, which ensured that the fewest possible number of channels were removed. After this process the number of remaining units in Monkey Pe was 78.3 ± 7.8 and 79.5 ± 8.5 (mean ± standard deviation), in Monkey Wa was 85.3 ± 8.1 and 25.2 ± 4.4, and in Monkey Sa was 62.6 ± 9.8 and 75.3 ± 13.2 for the right and left hemispheres respectively.

After the above preprocessing steps, the spiking activity was binned into spike counts. The bin was defined as the 1 second of the delay period immediately preceding the earliest possible saccade initiation cue in the session, such that each trial contributed one bin. For example, if the earliest possible saccade initiation cue in a given session was 1.5s after the target disappeared, the bin would be defined as 0.5-1.5s after the target disappeared for every trial in that session.

To increase the number of observations for model fitting, we combined trials across conditions (saccade target direction). To accomplish this, before fitting the pCCA-FA model or before computing *r*_*sc*_, we first subtracted the condition mean from the binned spike counts in each condition and then combined residual spike counts across conditions. An alternative approach is to z-score spike counts in each condition before combining^52^, which yielded qualitatively similar *r*_sc_ results.

#### Separation of slow and fast components

In this work, we are interested in trial-to-trial variability of neural activity on a timescale of seconds. As we investigated interactions between left and right PFC, it soon became apparent that both areas also contained a component that varied slowly on a timescale of minutes to hours. While slow-timescale variability presents its own set of scientific questions (e.g., Cowley et al. ^45^), these fluctuations complicated our analyses by introducing dependencies between one trial and the next, violating the basic independence assumption of correlation analysis (including regression, Pearson correlation, pCCA, and pCCA-FA; Figure S1). Therefore, we removed the slow component from the neural activity (Figure S1), and focused all analyses in this work on faster-timescale trial-to-trial variability, except where noted in Figure S1. We computed the slow component for each session using a centered boxcar filter of length 25 trials, computed after removing target information as described above. We then subtracted this component from the residual spike counts to remove slow-timescale trends that could have induced spurious correlations^50^. We performed this preprocessing procedure independently for each neuron. All analyses in this study were performed on the residual faster-timescale component.

#### Preprocessing of pupil diameter

Eye position and pupil diameter were monitored using monocular infrared tracking at a 1000 Hz sample rate (EyeLink 1000, RS Research, Mississauga, Canada). We measured pupil diameter in the time window starting 200ms before target onset until the saccade initiation cue on every trial. Each trial was smoothed using a median filter with a 50ms window (i.e., the median value in a 50ms centered window was subtracted from each sample) to remove high frequency noise. For the analyses in Figure 6B-C, we computed mean smoothed pupil diameter (arbitrary units) over the full time window described above. This yielded one average pupil diameter value per trial. Like the preprocessing of neural activity, we removed slow-timescale fluctuations in pupil diameter using a centered boxcar filter of length 25 trials. To ensure pupil diameter was on a similar scale across sessions, we z-scored the residual pupil diameter values for each session by subtracting the mean and dividing by the standard deviation of pupil diameter across the session.

For analyses in Figure 6D, we computed evoked pupillary responses. Evoked pupil diameter was defined as the change in pupil diameter following target onset. For each trial, we took the smoothed pupil diameter trace and measured the baseline before target onset as the average pupil diameter in a 100ms window ending 50ms before target onset (i.e., the window began 150ms before target onset). We then measured the average diameter in a 100ms window beginning 50ms after target onset and computed the difference between these two quantities for each trial. To compare across sessions, we z-scored the evoked pupil diameter within each session. As the evoked pupil diameter is already baseline-subtracted, no slow trends were removed in this analysis.

#### Probabilistic canonical correlation analysis

A commonly used approach for identifying interactions between brain areas is canonical correlation analysis (CCA)^32^. Briefly, CCA attempts to find a co-fluctuation pattern in area 1 and a co-fluctuation pattern in area 2, such that neural activity along these co-fluctuation patterns is maximally correlated across areas. Further co-fluctuation patterns can be found subject to the constraint that they are uncorrelated with previous co-fluctuation patterns. The correlations along these co-fluctuation patterns are known as canonical correlations (*ρ*). Note that CCA identifies activity shared across areas, but is not designed to partition activity into across- and within-area components.

The probabilistic interpretation of CCA (pCCA^62^) is defined by the following model. pCCA defines a linear-Gaussian relationship between the spike count vectors 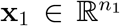 from area 1 and 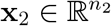 from area 2 and the across-area latent variables **z** ∈ ℝ^*d*^ as:

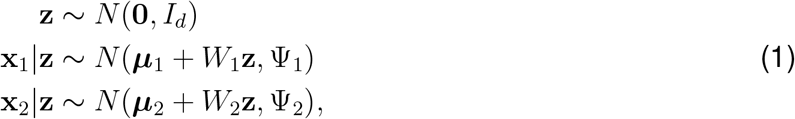

where *n*_1_ and *n*_2_ are the number of neurons recorded in areas 1 and 2, respectively, and *d* is the number of across-area latent variables. The identity matrix *I*_*d*_ ∈ ℝ^*d*×*d*^ describes prior covariance of the latent variables. The vectors 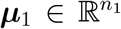 and 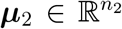 represent the mean spike counts of area 1 and area 2, respectively. The loading matrices 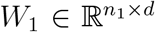 and 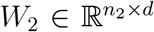 map the latent variables onto area 1 and area 2 respectively, and 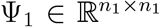 and 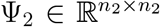 are full covariance matrices.

From Equation 1, the marginal distributions for **x**_1_ and **x**_2_ are:

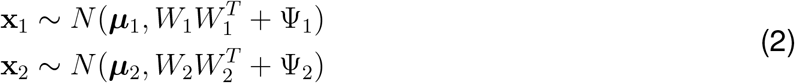

We can see that pCCA decomposes the covariance of neural activity from each area into a lowrank component shared across areas (e.g., 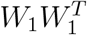) and a full “noise” covariance matrix (e.g., Ψ_1_).

Note that pCCA treats all variance which cannot be explained by across-area latent variables (**z**), including within-area interactions and variance independent to each neuron, as “noise”.

It can be shown that pCCA (Equation 1) finds the same co-fluctuation patterns (described by *W*_1_ and *W*_2_) as CCA that maximize correlation across areas^62^. CCA and pCCA therefore identify the same canonical correlations.

### Probabilistic canonical correlation analysis - factor analysis (pCCA-FA)

We developed a model called pCCA-FA (a combination of probabilistic canonical correlation analysis and factor analysis (FA^106^)), which explicitly partitions covariance of neural activity into an across-area component, a within-area component, and a component independent to each neuron. Though we term the components across-”area” and within-”area”, the pCCA-FA framework can be applied to any two simultaneously-recorded populations of neurons. pCCA-FA is related to extensions of FA that have been proposed in other contexts, such as Tucker ^107^ and Klami et al. ^108^.

pCCA-FA defines the linear-Gaussian relationship between the spike count vectors 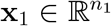 from area 1 and 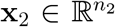 from area 2 and across-area latent variables **z** ∈ ℝ^*d*^ and within-area latent variables 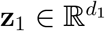 and 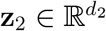 as:

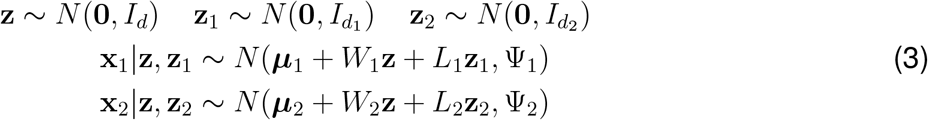

where *d* is the number of across-area latent variables, and *d*_1_ and *d*_2_ are the number of of withinarea latent variables for areas 1 and 2, respectively. The identity matrices *I*_*d*_ ∈ ℝ^*d*×*d*^, 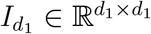, and 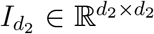 describe the prior covariance of each set of latent variables. Given *n*_1_ neurons from area 1, 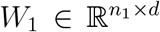 is the loading matrix for the across-area component in area 1, and 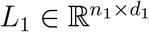 is the loading matrix for the within-area component in area 1. The matrix 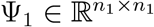 is a diagonal matrix containing the independent variance of each neuron in area 1, and 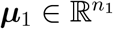 represents the mean spike counts of area 1. The parameters for area 2 are defined analogously.

From Equation 3, the marginal distributions for **x**_1_ and **x**_2_ are:

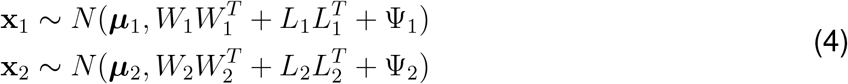

We can observe that pCCA-FA decomposes the covariance of neural activity in area 1 as the sum of an across-area component 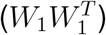, a within-area component 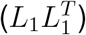, and an independent neuron component (Ψ_1_). The covariance of neural activity in area 2 is decomposed into analogous across-area 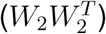, within-area 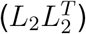, and independent neuron (Ψ_2_) components. In the pCCA-FA model, variance shared both with neurons in the same area and with neurons in the other area is assigned to across-area %sv (appearing in the pCCA-FA model as *W*_*m*_ for each area *m*, and corresponding to latent variable **z**, which involves both areas). Consequently, variance is assigned to the within-area component *only* when it is not shared with any neuron in the other area (appearing in the pCCA-FA model as *L*_*m*_, and corresponding to latent variable **z**_**m**_, which only involves area *m*).

pCCA-FA can be viewed as a combination of two existing latent variable methods, namely pCCA, which finds co-fluctuation patterns that maximize correlation between two populations, and FA, which maximizes covariance between neurons within a single population. Importantly, pCCA-FA explicitly identifies both types of covariance in a single probabilistic framework. The key difference between pCCA-FA and pCCA is that pCCA-FA constrains the full covariance matrices identified by pCCA (e.g., Ψ_1_ in Equation 2) as low-rank plus diagonal (e.g., 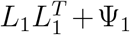 in Equation 4). It follows that pCCA-FA has fewer parameters than pCCA, and is thus more robust in the regime of low trial counts (i.e., abless to recover ground truth with fewer trials; Figure S3C-D). With sufficient trials, pCCA-FA and pCCA identify the same across-area co-fluctuation patterns.

### Fitting pCCA-FA and estimating latent variables

To fit pCCA-FA to neural activity and to estimate the latent variables, we define a joint vector of neural activity in both areas 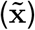 and a joint vector of across- and within-area latent variables 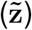:

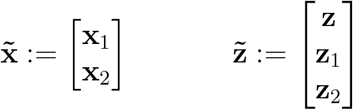

The probabilistic model of pCCA-FA from Equation 3 can be rewritten as:

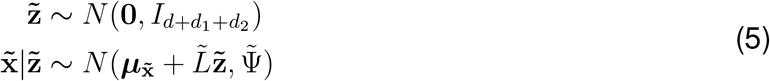

Where

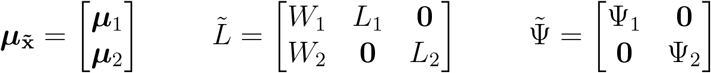

and ***µ***_1_, ***µ***_2_, *W*_1_, *W*_2_, *L*_1_, *L*_2_, Ψ_1_, and Ψ_2_ are defined as in Equation 3. The formulation in Equation 5 makes clear how we can think of pCCA-FA as a constrained and structured FA model. The zeros in the loading matrix 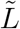 ensure that the within-area latent variables (**z**_1_ and **z**_2_) are only used to explain activity in their respective areas, whereas across-area latent variables (**z**) are used to explain activity in both areas. Based on Equation 5, the marginal distribution of the activity in both areas is:

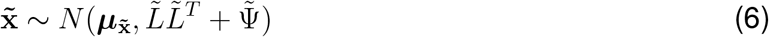

We fit pCCA-FA model parameters to neural activity using an exact expectation-maximization (EM) algorithm^109^. The EM algorithm for pCCA-FA resembles that used for FA. Specifically, the M-step update equations for the parameters 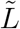 and 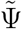 mirror those derived in FA for the loading matrix and noise covariance matrix. Each M-step parameter update for pCCA-FA (i.e., for parameters *W*_1_, *L*_1_, etc.) effectively applies a zeroing mask to the contents of 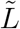 *L* ensuring that the blocked structure of the loading matrix is maintained. Additionally, there is no constraint on the relationship between the across- and within-area co-fluctuation patterns (e.g., they are not constrained to be orthogonal).

The columns of each loading matrix (*W*_*m*_ and *L*_*m*_ for each area *m*) as estimated using the EM algorithm are not ordered. As a post-processing step for all analyses, we applied the singular value decomposition to transform each loading matrix, such that the first column contained the co-fluctuation pattern that explained the most shared variance, the second column explained the second-most shared variance, etc. For across-area co-fluctuation patterns, it may also be desirable to order the columns according to shared correlation (as in CCA, see below).

For each session, we jointly chose the dimensionalities for across- (*d*) and within-area (*d*_1_ and *d*_2_) latent variables using 10-fold cross-validation and selected the dimensionalities which maximized the cross-validated data likelihood *P* 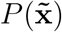 (see Equation 6). We tested integer dimensionality values between 1 and 15, inclusive, for each cross-validation run. After dimensionalities were selected, a final pCCA-FA model was fit with the specified dimensionalities using all trials. The parameters of this final model were used in all analyses.

To estimate the across- and within-area latent variables in pCCA-FA (e.g., for use in predicting pupil diameter in Figure 6), we computed the posterior mean of latent variables:

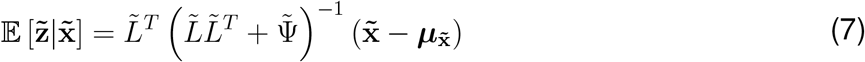

where the first *d* entries of 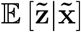 are 𝔼 [**z x**_1_, **x**_2_] (across-area latent variables), the next *d*_1_ entries are 𝔼 [**z**_1_ **x**_1_] (within-area latent variables for area 1), and the final *d*_2_ entries are 𝔼 [**z**_2_ **x**_2_] (withinarea latent variables for area 2).

### Computation of canonical variables

The pCCA-FA parameters allow for the computation of the canonical variables analogous to those found for CCA (Figure S6A). Following Bach and Jordan ^62^, the canonical directions and correlations can be obtained using singular value decomposition on the cross-correlation matrix *K* ∈ ℝ^*n*×*n*^:

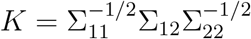

where *n* is the total number of neurons (i.e., *n*_1_ + *n*_2_),

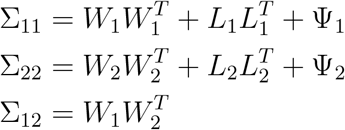

are the blockwise entries of 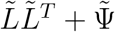. The canonical pairs of directions 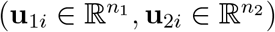 for *i* = 1, 2, …, *d*, are defined as:

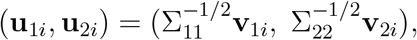

where 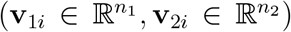 are a pair of left-right singular vectors of *K*. The corresponding singular value *ρ*_*i*_ is the canonical correlation.

Note that the Σ matrices are defined in terms of the estimated pCCA-FA parameters rather than the sample covariance matrices (which is what CCA and pCCA use). This is the regularization that pCCA-FA provides that enables it to perform well in a limited data regime (Figure S3C-D).

### Measuring percent shared variance (%sv) and dimensionality (*d*_*shared*_)

We defined two metrics to summarize the outputs of pCCA-FA: the percentage of shared variance (%sv) and shared dimensionality (*d*_*shared*_), which were originally introduced for FA^26,60^ and adapted here for pCCA-FA.

*%sv -* We used %sv to measure the prominence of (i.e., the proportion of variance explained by) the across- or within-area components. For a given pCCA-FA model, there are four instances of %sv: an across-area and a within-area instance, for each of the two areas. The across-area

%sv quantifies the percentage of each neuron’s variance that was explained by across-area latent variables (i.e., the percentage of each neuron’s activity that was shared with at least one other neuron in the other area). More specifically:

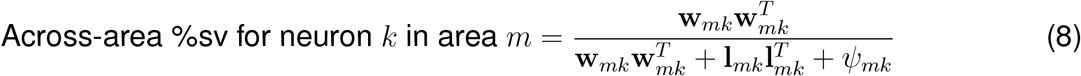

where **w**_*mk*_ is the *k*^*th*^ row of the across-area loading matrix for area *m*, **l**_*mk*_ is the *k*^*th*^ row of the within-area loading matrix for area *m*, and *ψ*_*mk*_ is the independent variance for the *k*^*th*^ neuron in area *m* (i.e., the *k*^*th*^ entry of the diagonal of Ψ_*m*_). The numerator represents the across-area variance for neuron *k* in area *m*, and the denominator represents the total variance for that neuron. In our analyses of %sv, we averaged over neurons in a brain area to obtain a single across-area %sv metric for each area.

Similarly, the within-area %sv quantifies the percentage of each neuron’s variance that was explained by within-area latent variables (i.e., the percentage of each neuron’s activity that was shared with at least one other neuron in the same area, but was not shared with any neurons in the other area). More specifically:

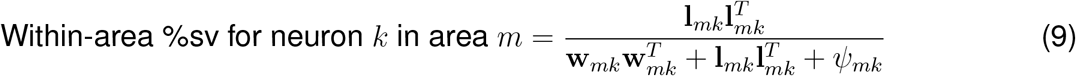

*d*_*shared*_ *-* We used the metric, *d*_*shared*_, to measure the “complexity” of across- and within-area interactions. For a given pCCA-FA model, there are four instances of *d*_*shared*_, an across-area and a within-area instance, for each of the two areas. Note that *d*_*shared*_ measures the dimensionality of interactions shared among neurons and excludes activity related to each neuron’s independent variance. We measured *d*_*shared*_ in an analogous way to Williamson et al. ^26^, but adapted for across- and within-area dimensionalities.

Briefly, the first step in measuring across-area *d*_*shared*_ is to find the number of across-area latent variables *d*^∗^ that maximizes the cross-validated data likelihood. We then fit the loading matrices *W*_*m*_ with *d*^∗^ across-area latent variables. Across-area *d*_*shared*_ is defined for each area *m* as the minimum number of latent variables needed to explain at least 95% of the across-area shared covariance in 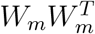, where *d*_*shared≤*_*d*^∗^. A 95% threshold was used in order to focus on latent variables that explain a large amount of shared variance. Note that although there is a single set of *d*^∗^ across-area latent variables, it is possible to obtain a different value of acrossarea *d*_*shared*_ for each area. We analogously defined within-area *d*_*shared*_ for each area using the procedure described above by replacing 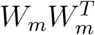 with 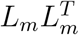.

### Simulated data generation

To validate the pCCA-FA model, as well as generate chance distributions (Figure 5D), we generated simulated data **x**_*m*_ for each area *m* from the pCCA-FA model according to Equation 4. This involved defining ground truth model parameters ***µ***_*m*_, *W*_*m*_, *L*_*m*_, and Ψ_*m*_ for each area *m*. For model validation, the ground truth parameters were defined as noted in the corresponding figure caption (Figures S3, S4). For Figure 5D, we defined these model parameters to be copies of those that were estimated from each of the neural recordings, with one manipulation, described below.

For the chance distributions in Figure 5D and the simulated data in Figure S4, we specified the angle between across- and within-area co-fluctuation patterns (*θ*_*sim*_). To achieve this, we held the across-area loading matrix (*W*_*m*_) constant and set the first column of the within-area loading matrix (*L*_*m*_) to be a vector that was *θ*_*sim*_ degrees apart from the first column of *W*_*m*_. The remaining columns of the within-area loading matrix were randomly generated from a standard normal distribution, and then orthogonalized relative to the first column (i.e., the first column remained fixed and the remaining columns were set to be mutually orthogonal to it).

### Pupil diameter regression

To explain pupil-related variance using the latent variables estimated by pCCA-FA, the across-area latent variables 𝔼 [**z**|**x**_1_, **x**_2_] and the within-area latent variables 𝔼 [**z**_1_|**x**_1_] and E [**z**_2_|**x**_2_] were estimated using Equation 7. We regressed either the across-area or the within-area (right or left PFC) latent variables onto the mean pupil diameter (Figure 6C) using a linear regression model for each set of latent variables. We then assessed *r*^2^, the proportion of variance in pupil diameter explained by the regression model.

To compute chance distributions, we repeated the procedure above, except that we performed the regression using latent variables estimated from the neural activity from a different session than the one in which we measured pupil diameter (i.e., the latent variables from session *i* were used to explain the pupil from session *j*, where *i* ≠ *j*). Trials were truncated in the session with more trials to ensure equal trial numbers, which resulted in chance distributions with 240 samples for Monkeys Pe and Sa with 16 sessions, and 90 samples for Monkey Wa with 10 sessions. We report the 95% confidence intervals of this chance distribution (Figure 6C, gray bars, termed “shuffle”). We performed regression for the pupillary evoked response (Figure 6D) in an analogous way.

To control for differences in across- and within-area dimensionality, we repeated the entire procedure using only one predictor (i.e., the top across- or within-area latent variable) rather than the full set of predictors (i.e., *d* across- or *d*_*m*_ within-area latent variables). This yielded similar results.

### sQuantification and statistical analysis

Analyses were conducted using custom code written in MATLAB (Mathworks, Natick, MA) and Python. All statistical details for relevant comparisons can be found in the corresponding figure captions. Significance was determined using a p-value of 0.05, and all p-values are directly reported. Strategies for inclusion of data are described in methods section “Preprocessing of neural activity”.

