## Supplemental Figures for "Interactions across hemispheres in prefrontal cortex reflect global cognitive processing"

**Supplemental information**

### Supplement inventory


- Figure S1
- Figure S2
- Figure S3
- Figure S4
- Figure S5
- Figure S6
- Supplemental references

### Supplemental figures

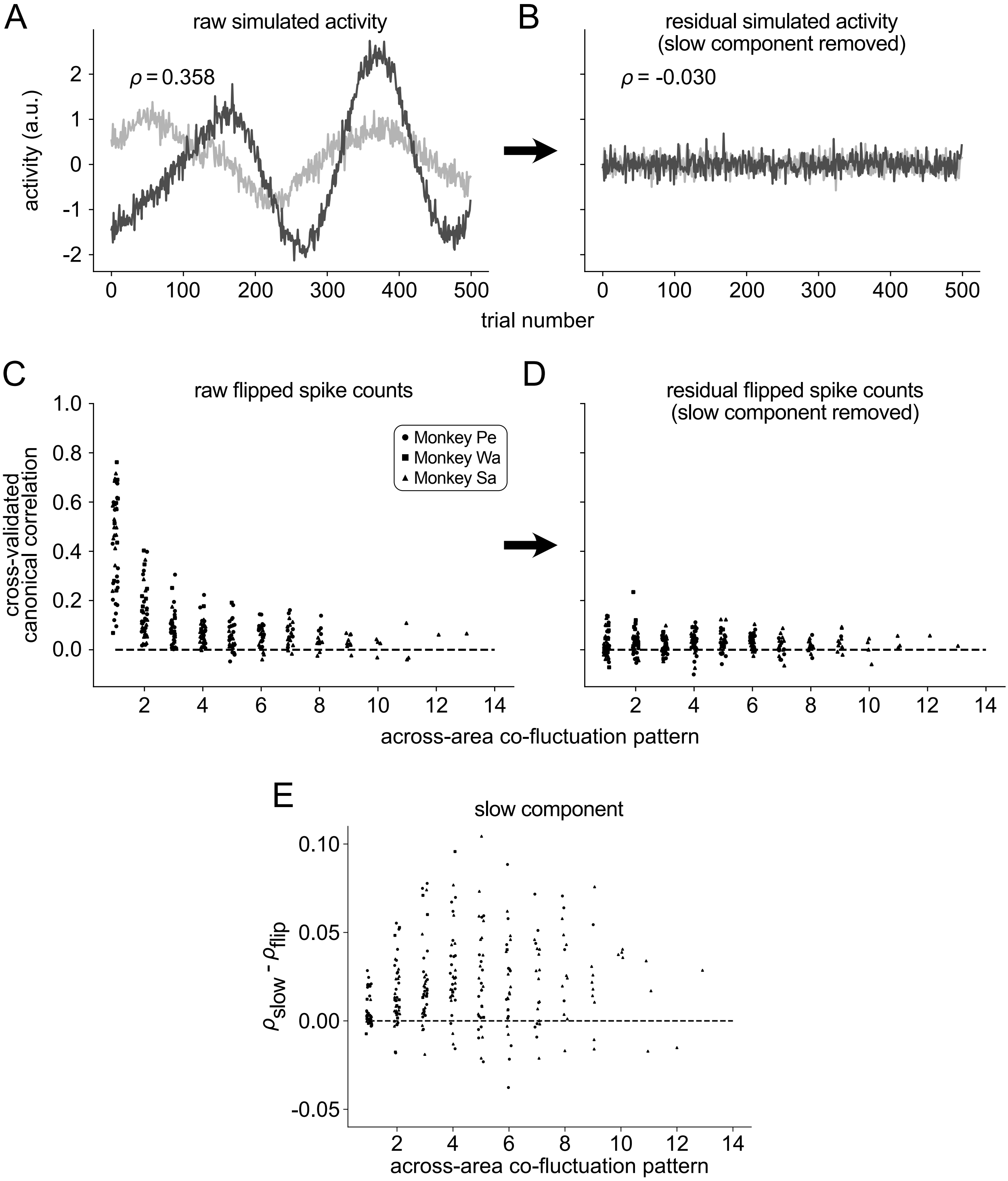


#### Figure S1. Spurious correlations induced by slow-timescale fluctuations, related to Figures 2 and 4, and STAR Methods.

In this work, we were interested in characterizing trial-to-trial variability shared among neurons. However, neural activity can exhibit multiple timescales of variability that can influence interpretation. For example, fluctuations on the timescale of minutes to hours have been observed in many brain areas, including PFC [S1]. It is important to control for these slow timescales in correlation analyses such as $r_{\text{sc}}$ and canonical correlation analysis (CCA) to prevent the measurement of potentially spurious correlations [S2, S3].

(A) To illustrate how slow-timescale activity fluctuations can yield spurious correlations, consider a simulation where activity from two brain areas is generated as two independently drawn Gaussian Processes (GPs, [S4]). Each GP consists of a time series of 500 values, representing the activity of a single brain area over 500 trials. We generated two GPs representing two brain areas with corresponding trial indices, each including a slow-timescale fluctuation with time constant $\tau = 50$ trials. Since the GPs were independently drawn, we would expect them to have a correlation across trials that is close to zero. However, given a limited number of trials, two independent GPs with slow-timescale fluctuations can have substantial non-zero correlation (here, the two time series have a trial-to-trial correlation of $\rho = 0.358$).

(B) As the number of trials grows to infinity, the correlation between the GPs would shrink to zero. However, this trial count inflation is not feasible in animal experiments. Instead, we can remove the slow-timescale components by subtracting a moving average (i.e., retain only the residual fast component; see STAR Methods). After this subtraction, the correlation between the two GPs is now close to zero ($\rho = -0.030$). Thus, whenever fluctuations in the activity occur on timescales that extend across a substantial portion of the trials, spurious correlations can arise.

(C) The same phenomenon could be seen in our neural recordings. We ran a control analysis to demonstrate the importance of removing slow components before performing any correlation analysis in neural activity. To generate a chance distribution, we performed a “flip control” where the trial order of neural activity in one area (right PFC) was flipped. Our assumption was that this flipping would break any trial-to-trial correspondence and should in principle result in correlations close to zero. Any recovered correlations would be spurious (induced by slow-timescale fluctuations).

For each experimental session, we fit a pCCA-FA model to flipped neural activity (left PFC remained the same, right PFC was flipped). We selected the dimensionalities of each model to be the same as what was identified in the non-flipped neural activity (i.e., in the models fit for analysis in Figures 4-6). We then computed cross-validated canonical correlation by splitting the trials of flipped neural activity into training and testing folds. For each fold, we estimated pCCA-FA model parameters using the training trials. Then, we computed the held-out posterior means ($E\left[\mathbf{z}|\mathbf{x}_1\right]$ and $E\left[\mathbf{z}|\mathbf{x}_2\right]$, see STAR Methods and Equation 7) using the testing trials. Across folds, this yielded one posterior mean per brain area per trial. The cross-validated canonical correlation is the trial-by-trial correlation of the posterior means. Note that this is equivalent to canonical correlations that would be identified by CCA, when applied to the same data.

When fitting pCCA-FA to flipped neural activity, we recovered large cross-validated canonical correlations, indicating that the across-area variance of models fit to raw spike counts could be representing spurious correlations rather than true across-area interactions. Each plotted point indicates the cross-validated canonical correlation (i.e., the correlation between held-out posterior means) for a pair of across-area co-fluctuation patterns (one in each area) in a single session. The horizontal axis positions of each point are jittered slightly for visual clarity.

(D) If the slow component were inducing spurious correlations, we would expect that removing this component from neural activity would reduce the correlations seen in (C). To test this, we subtracted the slow component of neural activity from the raw activity using a moving average (see STAR Methods). When fit to the residual fast component, pCCA-FA recovered cross-validated canonical correlations close to zero, indicating that spurious correlations were successfully removed from the neural activity. Therefore, the analyses in this work ($r_{\mathrm{sc}}$ in Figure 2 and pCCA-FA in Figures 4-6) focus on the residual fast component. We used the same model model fitting procedure and computation of cross-validated canonical correlations as in (C), but on the residual flipped neural activity. Each point indicates cross-validated canonical correlation for a pair of across-area co-fluctuation patterns as in (C).

(E) Though this work focuses on trial-to-trial variability, we also wondered whether any meaningful (i.e., not spurious) correlations existed in the slow component of neural activity. We fit pCCA-FA to the estimated slow components of neural activity. To determine whether the estimated canonical correlations ($\rho_{slow}$) were meaningful, we defined a chance distribution using the correlations recovered when fitting pCCA-FA to the flipped slow component ($\rho_{flip}$), using the same flip control as in (C). The selection of dimensionalities was performed as in (C) for the two pCCA-FA models. Similar to the analysis in our previous work [S1], we found here that there were substantial correlations present in the slowly varying neural activity. $\rho_{slow}$ was above $\rho_{flip}$ ($\rho_{slow} - \rho_{flip} > 0$) across animals for most across-area co-fluctuation patterns (Monkey Pe: $p < 0.0001$, Monkey Wa: $p = 0.0010$, Monkey Sa: $p < 0.0001$; paired t-test). This indicates that slow-timescale across-area interactions do indeed exist in PFC activity (perhaps similar to the correlations observed in our previous analysis of V4 and PFC [S1]), in addition to the trial-to-trial effects that we focus on in this work.

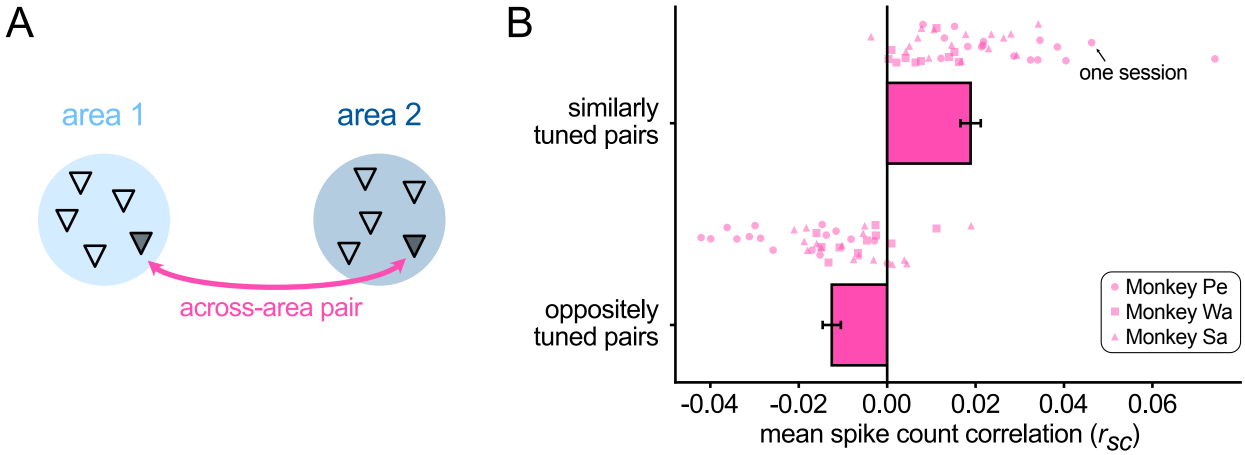


#### Figure S2. Relationship between noise correlation ($r_{\mathrm{sc}}$) and signal tuning in across-area neuron pairs, related to Figure 2

(A) In Figure 2C-D (bottom), we observed that the noise correlations ($r_{\mathrm{sc}}$) averaged over all across-area neuron pairs were near zero. Given this, one might have concluded that there were no across-area interactions in PFC.

(B) Here, we investigated if the distribution of across-area $r_{\mathrm{sc}}$, despite having a mean near zero, contained pairs of neurons with positive $r_{\mathrm{sc}}$ and other pairs of neurons with negative $r_{\mathrm{sc}}$, in way that was systematically related to the neurons' task tuning. To do this, we identified each neuron's tuning across all of the saccade target locations. We determined whether each pair of neurons had similar or opposite tuning by computing their signal correlation. This was defined as the Pearson correlation between the two neurons' trial-averaged responses to each of the possible targets (4, 8, or 16 depending on the session) during the 1 second window of the delay period immediately preceding the earliest possible saccade initiation cue (i.e., the same window used to compute $r_{\mathrm{sc}}$ in Figure 2). To assess significance for each neuron pair, we generated a chance distribution of correlation values by using a permutation test (shuffling the target location labels). We then labeled a neuron pair as having significant signal correlation (positive for similarly tuned and negative for oppositely tuned pairs) if their signal correlation was more extreme than the $99^{th}$ percentile of the chance distribution. Neuron pairs not meeting this criterion were excluded from this analysis.

We computed the mean noise correlation among these subsets of neuron pairs (similarly tuned and oppositely tuned) in each session. Comparing the mean $r_{\mathrm{sc}}$ values across sessions, we found that similarly tuned pairs of neurons exhibited positive noise correlation (top; mean $r_{\mathrm{sc}} = 0.019$; $r_{\mathrm{sc}} > 0, p < 0.0001$, t-test), whereas oppositely tuned pairs of neurons exhibited negative noise correlation (bottom; mean $r_{\mathrm{sc}} = -0.013$; $r_{\mathrm{sc}} < 0, p < 0.0001$, t-test). This did not have to be the case, as we defined subsets of pairs of neurons using their *signal* correlation, then evaluated their *noise* correlation. Thus, by incorporating information about the neurons' task tuning, we uncovered a richer correlation structure among across-area neuron pairs than what was suggested by than mean $r_{\mathrm{sc}}$ of all across-area neuron pairs.

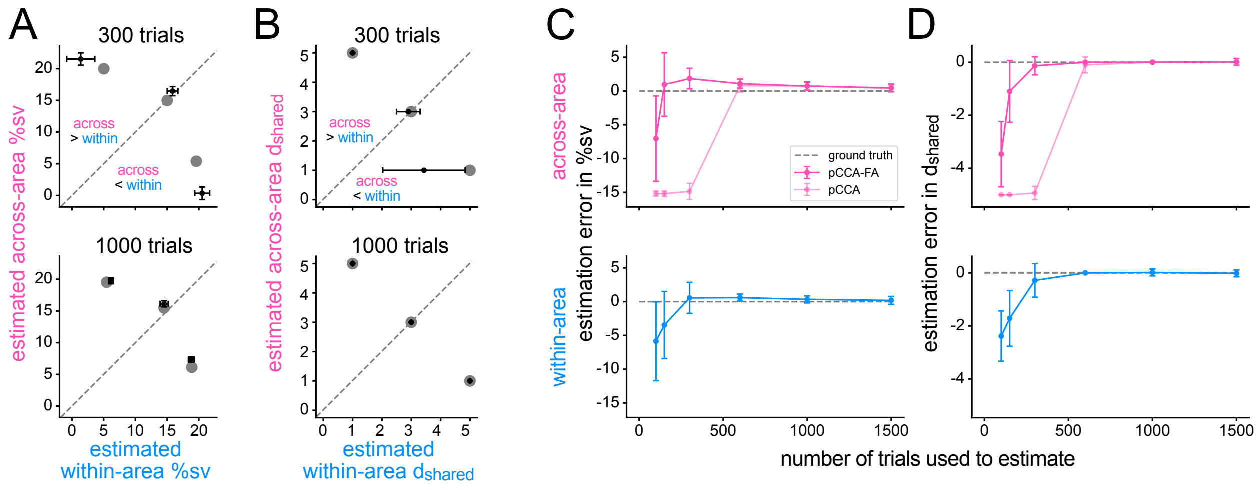


#### Figure S3. pCCA-FA accurately recovered %sv and $d_{shared}$ in simulations, related to Figure 3.

To validate the model fitting procedure, we asked whether pCCA-FA could recover ground truth values of %sv and $d_{shared}$ using realistic numbers of simulated neurons and trials. We systematically varied the ground truth values of across- and within-area %sv and $d_{shared}$ over values in the real-data regime. In each simulation, we used 30 neurons per area and generated random loading matrices $W_m$ and $L_m$ for each area $m$ using independent draws from a standard normal distribution. To scale the loading matrices to enforce a particular level of %sv, we adjusted the singular values of the across- and within-area loading matrices ($W_m$ and $L_m$) relative to each other to achieve the desired ratios. To facilitate this, we first applied the singular value decomposition to each loading matrix separately, which yielded orthogonal columns. In this analysis, we used a flat distribution of singular values for each loading matrix. Although this is not physiologically realistic (see Figure S6A-B), it was appropriate for this analysis to ensure that each co-fluctuation pattern described sufficient variance in cases of low %sv. The independent variance values (i.e., entries of $\Psi_m$) were then scaled to achieve a level of independent variance similar to what is estimated from our neural recordings (in this analysis, $75\%$). Across- and within-area latent dimensionalities used to calculate %sv were estimated in a cross-validated manner, as described in STAR Methods. We generated 30 independent datasets per parameter configuration, and error bars indicate one standard deviation computed across the 30 datasets.

(A) We tested three simulated %sv scenarios (marked with gray points), in which the across-area %sv was less than (below the dashed line), equal to (on the dashed line), or greater than (above the dashed line) the within-area %sv. This allowed us to assess whether pCCA-FA was able to accurately recover the across- and within-area components in scenarios in which one component overwhelmingly masked the other. To understand how trial count limitations impacted the estimation of %sv, we fit pCCA-FA to the three %sv scenarios using either 300 (top) or 1000 (bottom) trials. These trial counts represented the lower and upper end of the number of trials in our experimental sessions. Regardless of the relative proportions of across- and within-area %sv, we found that pCCA-FA was able to consistently identify the ground truth %sv (estimated %sv values near gray points indicating the ground truth configurations). With a small number of trials (top), when %sv was low pCCA-FA tended to underestimate the value of %sv because dimensionality was also underestimated. With sufficient trial counts (bottom), the estimated values of %sv became accurate (estimates on gray points with low spread). Each simulated dataset included 5 across-area latent variables and 3 within-area latent variables per area. These values were chosen to be within the range observed in our neural recordings. For visual clarity, we plotted the estimated %sv values from only one simulated brain area, but both areas yielded similar results.

(B) We next asked if pCCA-FA was similarly able to identify the ground truth $d_{shared}$ under scenarios in which across-area $d_{shared}$ was less than (below the dashed line), equal to (on the dashed line), or larger than (above the dashed line) within-area $d_{shared}$. We manipulated $d_{shared}$ while keeping the %sv constant ($\approx 15\%$ across-area and $\approx 10\%$ within-area). In all cases, pCCA-FA accurately estimated the across-area $d_{shared}$. When fitting pCCA-FA with 300 trials (top), we found that pCCA-FA underestimated within-area $d_{shared}$ in some cases. These were cases in which each within-area co-fluctuation pattern explained only $2 - 3\%$ of the overall variance, making them difficult to identify with limited trials. With increased trial counts (1000, bottom), pCCA-FA made no errors in estimating across- or within-area $d_{shared}$. As in (A), we plotted the estimated $d_{shared}$ values from only one area, but both areas yielded similar results. Taken together, these results demonstrate that pCCA-FA was able to identify and properly partition across- and within-area shared variability even with modest trial counts.

(C) For the simulated datasets in (A-B), we used either 300 or 1000 trials to fit each pCCA-FA model. Here, we systematically asked how the accuracy of the estimated %sv depended on the number of trials used to fit the model. The simulated datasets shown here each had 5 across-area latent variables with %sv $\approx 15\%$, and 3 within-area latent variables per area with %sv $\approx 10\%$. These values are within the ranges of those tested in (A-B). For each trial count configuration, we generated 30 sets of ground truth parameters and a single simulated dataset per set of parameters. We varied the number of trials used to fit pCCA-FA, and found that the estimated across- (top) and within-area (bottom) %sv values approached the ground truth value as the number of trials increased, in that the estimation errors decreased to zero.

(D) We next asked how the accuracy of the estimated $d_{shared}$ depended on the number of trials. At low trial counts, both across- (top) and within-area (bottom) $d_{shared}$ were underestimated. This likely caused the underestimation of %sv when trial counts were low. Estimates of $d_{shared}$ converged on the ground truth values as the trial count increased. Note that the simulations in (C-D) are paired (i.e., the models for 100 trials in (C) are the same as in (D)).

(C-D) A widely-used alternative and simpler model for identifying across-area variance is CCA or its probabilistic variant pCCA [S5]. Here we wondered whether applying pCCA-FA affords any advantages over pCCA in the real-data regime. Mathematically, pCCA does not separate within-area shared variance from variance independent to each neuron. Instead, it estimates a full “noise” covariance matrix for each area (see Equation 2) that requires more parameters than pCCA-FA (see Equation 4). Therefore, we expected that pCCA-FA would be able to accurately estimate %sv (C) and $d_{shared}$ (D) with fewer trials than pCCA.

To assess this, we applied pCCA to the same sets of simulated data as pCCA-FA. At low trial counts, we found that pCCA, like pCCA-FA, underestimated across-area %sv (C) because dimensionality was underestimated (D). However, while pCCA-FA’s estimate of across-area %sv quickly improved with more trials, pCCA’s estimate of across-area %sv required a greater number of trials to become accurate. Similar to %sv, we found that pCCA underestimated across-area $d_{shared}$ when trial counts were low, and required more trials than pCCA-FA to achieve accurate estimation. In practice, the lower number of parameters in the pCCA-FA model provides great benefit over pCCA in the regime of low to modest trial counts ($< 600$ trials). Both models achieved accurate estimates of across-area %sv and $d_{shared}$ in the tested configurations with sufficient trial counts ($> 600$ trials). In the regime of low trial counts, pCCA-FA and pCCA do not identify the same across-area co-fluctuation patterns, with the estimates from pCCA-FA being more similar to the ground truth co-fluctuation patterns. pCCA-FA and pCCA converge to identify identical across-area co-fluctuation patterns, which are also obtained by applying CCA to the same data [S5]. Note that pCCA is not designed to separate within-area and independent variability, and thus we could not assess its ability to identify within-area %sv or $d_{shared}$.

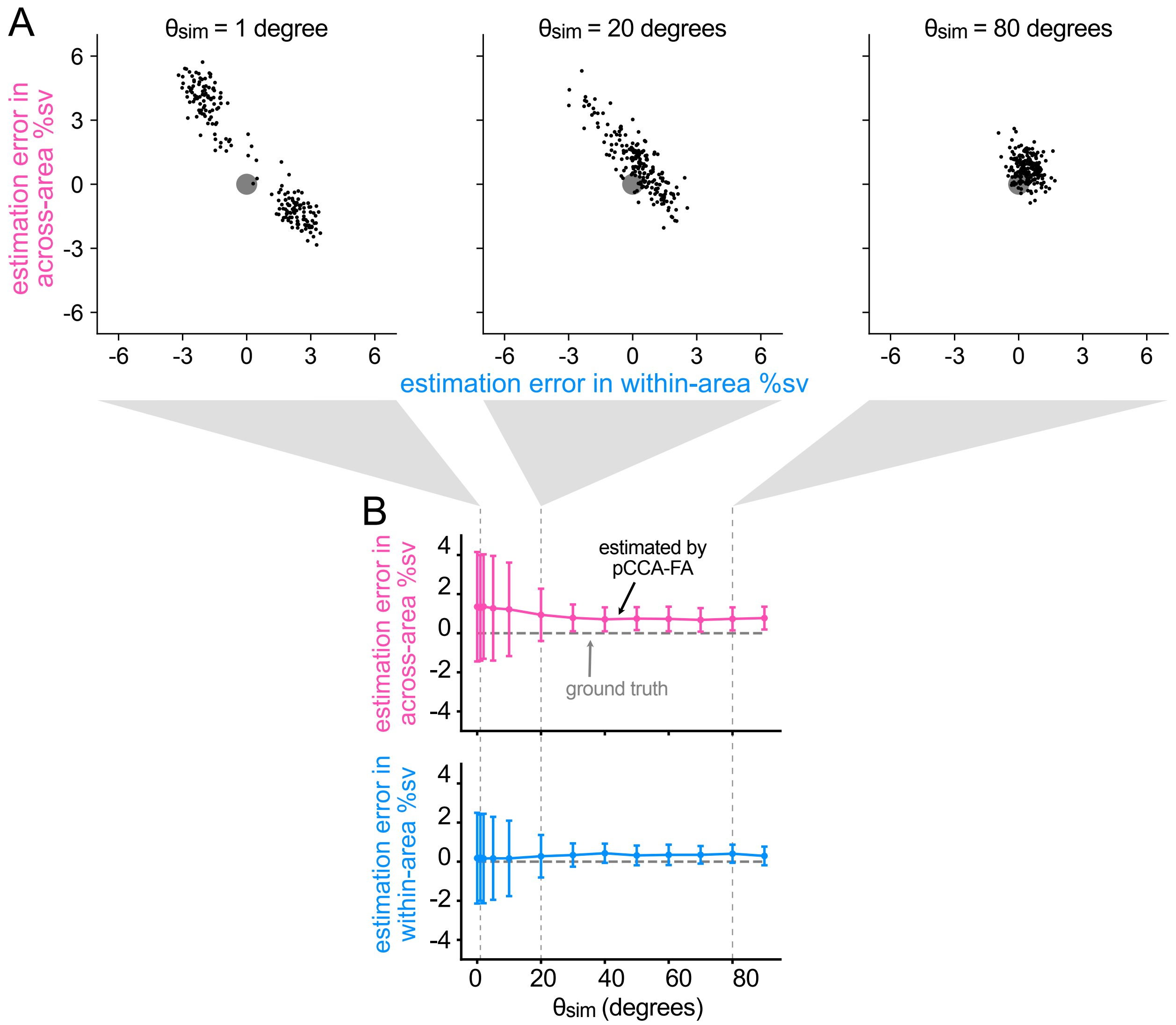


**Figure S4. Recovery of %sv depends on angle between across- and within-area co-fluctuation patterns, related to Figures 3 and 5.**

In Figure 5, we found that the angle between across- and within-area co-fluctuation patterns tended to be large (between $60$ and $90°$). We asked how well pCCA-FA could partition across- and within-area variance as the across- and within-area co-fluctuation patterns became more similar to each other. To test this, we conducted simulations in which we systematically varied the angle between the ground truth across- and within-area co-fluctuation patterns. We then asked how accurately pCCA-FA estimated across- and within-area %sv. In each simulation, we specified the angle between the top across- and within-area co-fluctuation pattern (i.e., the angle between the first column of the across- and within-area loading matrices; $\theta_{sim}$, see STAR Methods). We wished to test the dependence of the estimation procedure only on the value of $\theta_{sim}$, so we matched the level of %sv across all simulations and all values of $\theta_{sim}$. To do this, we generated loading matrices $W_m$ and $L_m$ for each area $m$ as in Figure S3, and scaled their columns to ensure that the ground truth value of across-area %sv was $\approx 15\%$ and within-area %sv was $\approx 10\%$. In this analysis, we scaled the columns of each loading matrix using singular values that followed an exponential distribution (i.e., the first co-fluctuation pattern had the highest singular value, and each subsequent co-fluctuation pattern had smaller singular values). This ensured that application of the singular value decomposition did not change the order of the columns, which would lead to an incorrect value of $\theta_{sim}$. The independent variance values (i.e., entries of $\Psi_m$) were then scaled to achieve $75\%$ independent variance. We selected other parameters to be within the range of values observed in neural recordings: 30 neurons per area, 5 across-area latent variables and 3 within-area latent variables for each area. We repeated the simulation procedure 100 times for each value of $\theta_{sim}$, using 1000 trials per simulated dataset. In this analysis, we were interested in the partitioning of across- and within-area shared variance, so we fit a single model using all trials at the ground truth values of across- and within-area dimensionality, then computed the estimated values of across- and within-area %sv. We report the error in estimated across- or within-area %sv which was computed as (estimated - ground truth).

(A) When the across- and within-area co-fluctuation patterns were aligned ($\theta_{sim} = 1°$; left), %sv was within $\approx 5\%$ of the ground truth value (black dots indicating each simulated population away from gray dot indicating ground truth of zero error). Each individual simulation contributes two points to the scatter, one per simulated population. As the co-fluctuation patterns became more distinct ($\theta_{sim} = 20°, 80°$; middle and right, respectively), pCCA-FA became more accurate in estimating across- and within-area %sv (black points overlapping gray point).

(B) To provide a fuller picture of the dependence of estimated %sv on $\theta_{sim}$, we systematically varied the alignment of the across- and within-area co-fluctuation patterns ($\theta_{sim}$). We then asked how the estimated partitioning of variance depended on the alignment of the co-fluctuation patterns. Estimated %sv settled to the ground truth value and variance across estimates decreased quickly as co-fluctuation patterns became more distinct (from left to right). Note that the results shown at $\theta_{sim} \in [1°, 20°, 80°]$ (marked by dashed lines) match the panels in (A). The error bars indicate the mean and standard deviation across 100 independent simulations (200 values; one per simulated brain area, yielding two per simulation).

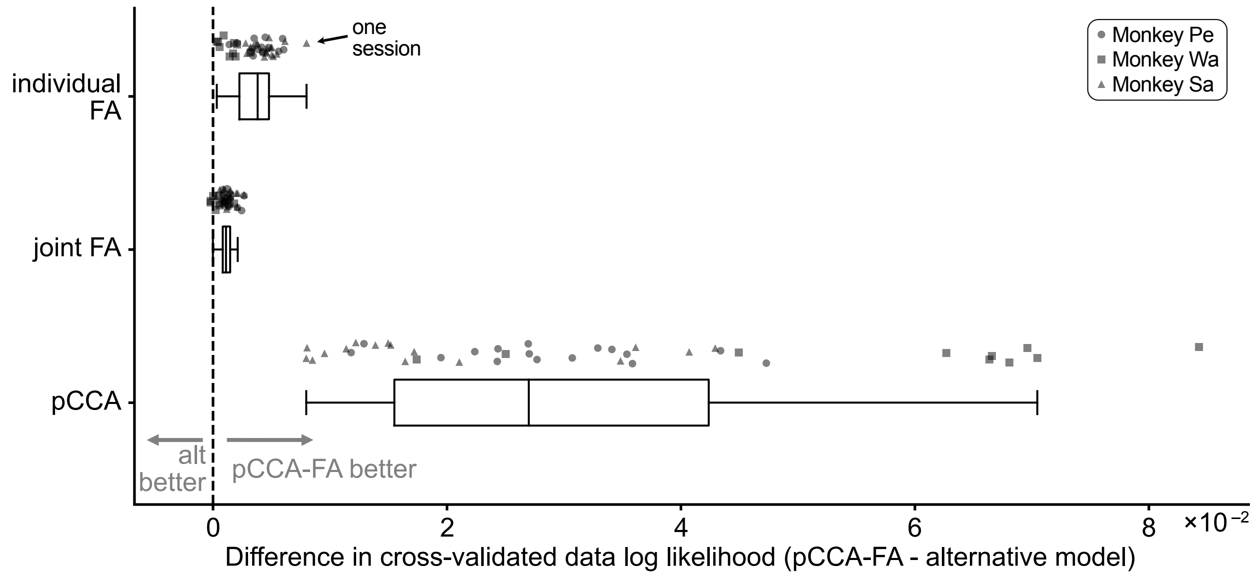


#### Figure S5. pCCA-FA outperformed alternative models on neural recordings, related to Figure 3.

Here we compared pCCA-FA to alternative statistical models that can be used to describe neural activity recorded simultaneously from two areas. We focused on probabilistic models tailored for studying trial-to-trial variability (without a time series model). Overall, we found that pCCA-FA outperformed alternative models on our dual hemisphere PFC recordings (measured in terms of higher cross-validated data likelihood). In addition to the greater generalization performance, pCCA-FA distinguishes between across- and within-area interactions, providing a more thorough dissection of neural activity than the alternative models that we considered.

For each recording session and model, we performed 10-fold cross-validation to identify the latent dimensionalities, and used the cross-validated data log likelihood of the selected model for comparison with other models (see Equation 6 in STAR Methods for the pCCA-FA definition). For each experimental session, we computed the difference in data log likelihood between pCCA-FA and that of each alternative model under consideration. A value above zero indicates that pCCA-FA outperformed the alternative model. For plotting purposes, we normalized the log likelihood of each session by the number of trials and neurons so that the log likelihood values would be comparable across sessions. Paired statistical tests (see below) were performed using non-normalized values. Each box plot covers the first to third quartiles of the differences in log likelihood, with the median marked with a line.

(top) For two populations of neural activity, we can fit a separate FA model to each area (termed “individual FA” model). Although this model does not identify across-area interactions, it is a simple model whose performance served as a baseline for interpreting the performance of pCCA-FA and other models. Note that this model is mathematically equivalent to pCCA-FA when the across-area dimensionality is set to 0. We fit a FA model to each area, and calculated the data log likelihood for both areas by adding together the log likelihood from each single-area model. We found that pCCA-FA outperformed the individual FA model (one point per session; Monkey Pe: $p < 0.0001$, Monkey Wa: $p = 0.0018$, Monkey Sa: $p < 0.0001$; paired t-test) because pCCA-FA captures activity that is correlated across areas, whereas the individual FA model does not.

(middle) We next fit FA jointly to all recorded neurons, without consideration that they were recorded from different areas (termed “joint FA” model). The joint FA model captures both across-

and within-area interactions together, and is not designed to separate these two types of interactions as pCCA-FA is. Again, we found that pCCA-FA outperformed the joint FA model (Monkey Pe: $p < 0.0001$, Monkey Wa: $p = 0.014$, Monkey Sa: $p < 0.0001$; paired t-test).

(bottom) Both formulations of FA considered so far are unable to identify across-area interactions separately from within-area interactions. To explicitly identify across-area interactions, we fit a pCCA model (see STAR Methods, [S5]) to the two areas. We found that pCCA-FA outperformed pCCA for all sessions (Monkey Pe: $p < 0.0001$, Monkey Wa: $p < 0.0001$, Monkey Sa: $p < 0.0001$; paired t-test). There were $\approx 450 - 850$ trials in each session, which is in the regime in which pCCA-FA outperformed pCCA in simulated data (Figure S3C-D). pCCA-FA outperforms pCCA in this regime because pCCA-FA has fewer parameters than pCCA (see STAR Methods).

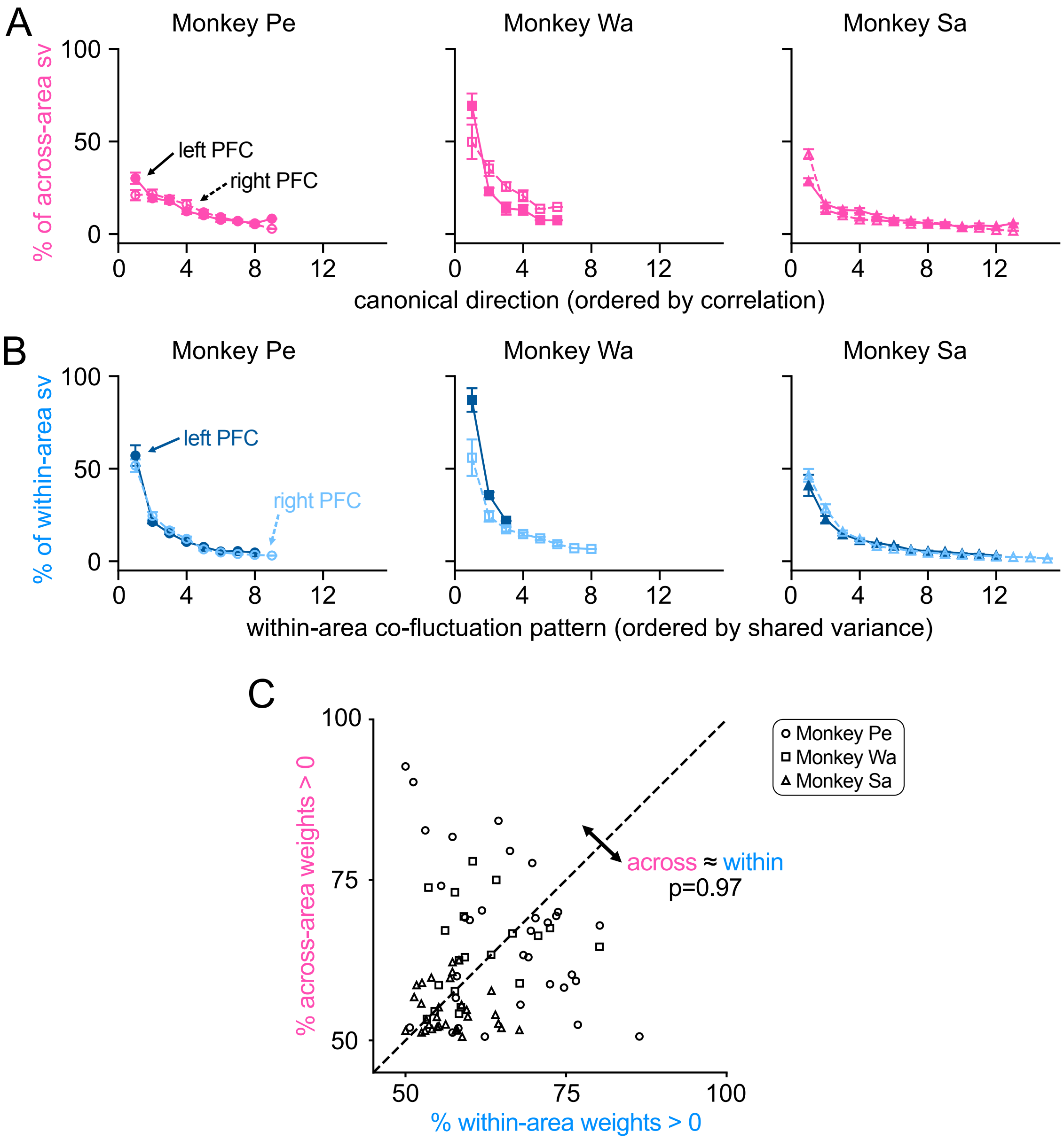


#### Figure S6. Characterizing across- and within-area co-fluctuation patterns, related to Figure 5.

In Figure 5 we compared the similarity of the across- and within-area co-fluctuation patterns identified by pCCA-FA. Here, we analyze the relative strength (A-B) and the diversity of weights onto the neural populations (C) of the co-fluctuation patterns.

(A) We first wondered how dominant the first co-fluctuation pattern of each type was relative to the subsequent co-fluctuation patterns. The across-area latent variables in pCCA-FA are defined by the co-fluctuations pattern each area that explain the most across-area shared variance. The pCCA-FA model also provides the capability to identify the most correlated co-fluctuation patterns across areas (as in CCA and pCCA). Here, we sought to understand whether the co-fluctuation patterns that described the greatest correlation across areas also explained the most shared variance across areas. We computed the proportion of across-area shared variance explained by each of the most correlated co-fluctuation patterns across areas. These co-fluctuation patterns identified by pCCA-FA correspond to the canonical directions identified by CCA or pCCA (see STAR Methods). We found that the co-fluctuation patterns that described the greatest correlation across areas indeed also tended to explain the most variance.

Specifically, we computed the percentage of across-area shared variance explained by the $i^{th}$ canonical direction $\mathbf{u}_{mi} \in \mathbb{R}^{n_m}$ in area $m$ as: $\frac{\mathbf{u}_{mi}^T(W_m W_m^T)\mathbf{u}_{mi}}{tr(U_m^T(W_m W_m^T)U_m)}$, where $tr(\cdot)$ is the trace, and $U_m \in \mathbb{R}^{n_m \times d}$ is the matrix whose columns contain the $d$ canonical directions $\mathbf{u}_{mi}, i = 1, 2, ..., d$ (see STAR Methods). Note that even though two curves are shown (one for neurons in right PFC, one for neurons in left PFC), the across-area variance is described by a single set of latent variables that involves both areas. We plotted the mean and standard error across sessions in each animal separately. As such, the plotted values need not sum to $100\%$ (though they do for each individual session).

(B) We also assessed the relative strength of the within-area co-fluctuation patterns. By definition, the within-area latent variables identified by pCCA-FA explain the greatest shared variance within an area that is not explained by activity in the other area. As a post-processing step, we reorder the within-area co-fluctuation patterns by the amount of shared variance explained, such that the first co-fluctuation pattern explains the greatest shared variance, the second co-fluctuation pattern explains the second-most shared variance, etc. (see STAR Methods). The percentage of within-area shared variance explained by the $i^{th}$ within-area co-fluctuation pattern in area $m$ is computed as: $\frac{\mathbf{a}_{mi}^T L_m L_m^T \mathbf{a}_{mi}}{tr(L_m L_m^T)}$, where $\mathbf{a}_{mi}$ is the $i^{th}$ eigenvector of $L_m L_m^T$. The numerator of this expression is equivalent to the $i^{th}$ eigenvalue of $L_m L_m^T$. Note that the two curves shown here now represent two different sets of latent variables identified by the pCCA-FA model: one involving neurons in right PFC (light blue), and one involving neurons in left PFC (dark blue). Across sessions, we found that the top co-fluctuation pattern explained a large proportion of shared variance relative to the subsequent co-fluctuation patterns (sharp drop off from first to second co-fluctuation patterns). This indicated that much of the within-area shared variance could be attributed to a single mode of variance, a finding that is sometimes termed "low rank variability" [S6].

(C) In (A-B) we found that the top across- and within-area co-fluctuation patterns were dominant relative to the subsequent co-fluctuation patterns. We next examined the diversity of weights onto the neural populations of these top co-fluctuation patterns, which indicates the extent to which a co-fluctuation pattern involved all neurons increasing or decreasing their activity together. For example, if the across-area variance involved a global modulatory process, all neurons might increase and decrease their activity together. If this were the case we would observe many weights of the same sign, or a low diversity of weights.

We quantify the diversity of weights by computing the percentage of same-signed weights for the top across- and within-area co-fluctuation pattern in each area (a total of four values per session). This metric corresponds to the proportion of values of the top across- or within-area co-fluctuation pattern sharing the same sign. The overall signed orientation of a co-fluctuation pattern is arbitrary, as flipping the sign of every weight along a co-fluctuation pattern yields an equivalent model. We oriented the sign of each co-fluctuation pattern such that the majority of weights were positive (i.e., the minimum percentage of same-signed weights is 50%, which we notate as positive by convention). We performed the orientation change on each individual co-fluctuation pattern to ensure a fair comparison between across- and within-area co-fluctuation patterns. This sign change was only performed for this analysis. Each plotted point corresponds to the top across- and within-area co-fluctuation pattern from one area, such that each session contributes two points (one point for each area).

We found that neither across- nor within-area co-fluctuation patterns had weights that were close to 100% positive. Rather than all neurons increasing and decreasing their activity together, the across- and within-area co-fluctuation patterns had diverse weights onto the neural populations. In fact, the across- and within-area co-fluctuation exhibited similar diversity in their weights. There was no significant difference between the percentage of positive weights in the top across- and within-area co-fluctuation patterns ($p = 0.97$ pooled across animals; Monkey Pe: $p = 0.80$, Monkey Wa: $p = 0.19$, Monkey Sa: $p = 0.035$; paired t-test). Notably this analysis does not include slow-timescale fluctuations in neural activity (which were removed in our preprocessing, see STAR Methods), but instead focuses on fast-timescale trial-to-trial variability. Our previous work, across several brain areas and many animals [S1, S7, S8, S9], has investigated the diversity of weights identified from neural population activity including slow-timescale fluctuations and found varied effects.

[S2] Kenneth D. Harris. Nonsense correlations in neuroscience. Preprint at bioRxiv, 10.1101/2020.11.29.402719, June 2021.

[S3] Evren Gokcen, Anna I. Jasper, João D. Semedo, Amin Zandvakili, Adam Kohn, Christian K. Machens, and Byron M. Yu. Disentangling the flow of signals between populations of neurons. *Nature Computational Science*, 2(8):512–525, August 2022.

[S4] Carl Edward Rasmussen and Christopher K. I. Williams. *Gaussian Processes for Machine Learning*. The MIT Press, 2006.

[S5] Francis R Bach and Michael I Jordan. A Probabilistic Interpretation of Canonical Correlation Analysis. Technical Report 688, University of California, Berkeley, Department of Statistics, 2005.

[S6] Chengcheng Huang, Douglas A Ruff, Ryan Pyle, Robert Rosenbaum, Marlene R Cohen, and Brent Doiron. Circuit models of low-dimensional shared variability in cortical networks. *Neuron*, 101(2):337–348, 2019. Publisher: Elsevier.

[S7] Jay A. Hennig, Emily R. Oby, Matthew D. Golub, Lindsay A. Bahureksa, Patrick T. Sadtler, Kristin M. Quick, Stephen I. Ryu, Elizabeth C. Tyler-Kabara, Aaron P. Batista, Steven M. Chase, and Byron M. Yu. Learning is shaped by abrupt changes in neural engagement. *Nature Neuroscience*, 24(5):727–736, May 2021.

[S8] Richard Johnston, Adam C Snyder, Sanjeev B Khanna, Deepa Issar, and Matthew A Smith. The eyes reflect an internal cognitive state hidden in the population activity of cortical neurons. *Cerebral Cortex (New York, NY)*, 32(15):3331–3346, December 2021.

[S9] Richard Johnston and Matthew A. Smith. Brain-wide arousal signals are segregated from movement planning in the superior colliculus. *eLife*, 13, August 2024. Publisher: eLife Sciences Publications Limited.
